# CRISPR-inhibition screen for lncRNAs linked to melanoma growth and metastasis

**DOI:** 10.1101/2024.07.24.604899

**Authors:** Stavroula Petroulia, Kathryn Hockemeyer, Shashank Tiwari, Pietro Berico, Sama Shamloo, Seyedeh Elnaz Banijamali, Eleazar Vega-Saenz de Miera, Yixiao Gong, Palaniraja Thandapani, Eric Wang, Michael Schulz, Aristotelis Tsirigos, Iman Osman, Ioannis Aifantis, Jochen Imig

**Affiliations:** Chemical Genomics Centre of the Max Planck Society, 44227 Dortmund, Germany; Max Planck Institute of Molecular Physiology, 44227 Dortmund, Germany; Department of Pathology and Laura & Isaac Perlmutter Cancer Center, NYU School of Medicine, New York 10016, NY, USA; Laura and Isaac Perlmutter Cancer Center, New York University, School of Medicine, New York 10016, NY, USA; Applied Bioinformatics Laboratories, Office of Science and Research, New York University School of Medicine, New York 10016, NY, USA; Interdisciplinary Melanoma Cooperative Group (IMCG), NYU Perlmutter Cancer Center, NYU Langone Health, New York, NY, USA; Departments of Urology and Medicine, New York University School of Medicine, New York, NY, USA; The Ronald O. Perelman Department of Dermatology, New York University School of Medicine, New York, NY, USA; The Jackson Laboratory for Genomic Medicine, Farmington, CT, USA

**Keywords:** LncRNA, CRISPRi screen, melanoma, metastasis, high-throughput sequencing

## Abstract

Melanoma being one of the most common and deadliest skin cancers, has been rising since the past decade. Patients at advanced stages of the disease have very poor prognoses, as opposed to at the earlier stages. Nowadays the standard-of-care of advanced melanoma is resection followed by immune checkpoint inhibition based immunotherapy. However, a substantial proportion of patients either do not respond or develop resistances. This underscores a need for novel approaches and therapeutic targets as well as a better understanding of the mechanisms of melanoma pathogenesis. Long non-coding RNAs (lncRNAs) comprise a poorly characterized class of functional players and promising targets in promoting malignancy. Certain lncRNAs have been identified to play integral roles in melanoma progression and drug resistances, however systematic screens to uncover novel functional lncRNAs are scarce. Here, we profile differentially expressed lncRNAs in patient derived short-term metastatic cultures and BRAF-MEK-inhibition resistant cells. We conduct a focused growth-related CRISPR-inhibition screen of overexpressed lncRNAs, validate and functionally characterize lncRNA hits with respect to cellular growth, invasive capacities and apoptosis in vitro as well as the transcriptomic impact of our lead candidate the novel lncRNA XLOC_030781. In sum, we extend the current knowledge of ncRNAs and their potential relevance on melanoma.

**Significance:** Previously considered as transcriptional noise, lncRNAs have emerged as novel players in regulating many cellular aspects in health and disease including melanoma. However, the number and as well as the extent of functional significance of most lncRNAs remains elusive. We provide a comprehensive strategy to identify functionally relevant lncRNAs in melanoma by combining expression profiling with CRISPR-inhibition growths screens lowering the experimental effort. We also provide a larger resource of differentially expressed lncRNAs with potential implications in melanoma growth and invasion. Our results broaden the characterized of lncRNAs as potential targets for future therapeutic applications.

## 1. Introduction

Malignant melanoma of the skin represents a significant clinical challenge due to its aggressive nature and propensity for metastasis (Bhatia, Tykodi, & Thompson, 2009). Despite advances in therapeutic strategies such as targeted and immune therapies (Schadendorf et al., 2015), melanoma remains one of the deadliest forms of skin cancer, with rising incidence rates globally (Garbe & Leiter, 2009). The clinical management of melanoma necessitates a deeper understanding of its molecular underpinnings to improve diagnostic accuracy, prognosis, and therapeutic outcomes.

From that perspective, long non-coding RNAs (lncRNAs) have emerged as pivotal regulators of gene expression and cellular processes, orchestrating diverse biological functions (Geisler & Coller, 2013; Gil & Ulitsky, 2020) LncRNA dysregulation has been implicated in various pathological conditions, including melanoma (Leucci et al., 2016; Xiao, Xia, Wang, & Xue, 2023), in which regulatory roles of lncRNAs are increasingly recognized, however not fully understood. Moreover, the ability of melanoma cells to acquire invasive properties also by the involvement of lncRNAs (Damsky, Theodosakis, & Bosenberg, 2014; Flaherty, Hodi, & Fisher, 2012; Xiao et al., 2023) underscores the importance of identifying novel therapeutic targets and predictive biomarkers. LncRNAs exert regulatory effects on gene expression at multiple levels, including chromatin remodeling, transcriptional regulation, and post-transcriptional processing, thus modulating key signaling pathways implicated in melanoma pathogenesis (Xiao et al., 2023). Several lncRNAs have been involved in melanoma development, progression, and therapeutic response. Notably, dysregulated expression of lncRNAs such as MALAT1, HOTAIR, and SAMMSON has been associated with melanoma metastasis, invasion, and resistance to chemotherapy (Leucci et al., 2016; Tang, Zhang, Su, & Yu, 2013; Tian, Zhang, Hao, Fang, & He, 2014) These lncRNAs function as oncogenic drivers by promoting tumor cell proliferation, migration, and epithelial-mesenchymal transition (EMT), however, our common understanding remains far from complete. Overall, elucidating the intricate interplay between melanoma and lncRNAs holds great promise for advancing our understanding of disease pathogenesis and improving clinical outcomes. Thus, improved understanding of how lncRNAs regulate melanoma pathogenesis will offer novel insights into mechanisms of progression and therapeutic resistance in this disease.

In the present work we aimed to define the extent to which lncRNAs contribute to melanoma progression by conducting a selective and integrative screening approach. We systematically executed a series of multi-step filtering selection of lncRNAs in malignant melanoma including: **1)** pre-identification of overexpressed lncRNAs in clinically relevant metastatic melanoma against healthy melanocytic samples as well as multi-drug resistant vs. parental cell lines hypothesizing an oncogenic gain-of-function thereof. **2)** a focused CRISPR-inhibition lncRNA growth screen in melanoma cells to unambiguously identify candidates which affect cell survival and fitness. We leveraged this strategy to gain the two-fold benefit of increasing the screening hit rate while lowering the effort and screening cost as whole lncRNA transcriptome CRISPRi analyses are challenging to conduct. Lastly, we validate and functionally characterize individual lncRNAs for growth and invasion related impact *in cellulo*.

## 2. Methods

### 2.1. Cell Culture

Lenti-X™ 293T cells were purchased from Takara Bio. 501-mel and Lenti-X™ 293T were grown in Gibco Dulbecco’s modified Eagle’s medium (DMEM), high glucose with pyruvate supplemented with 10% (v/v) fetal bovine serum (FBS) and 1% (v/v) penicillin/streptomycin. WM1361A were cultured in medium containing 80% (v/v) MCDB153 with trace elements, L-glutamine and 28 mM HEPES, 20% (v/v) Leibovitz L-15, and supplemented with 1.2 g/L NaHCO3, 2% heat inactivated FBS, 1.68 mM CaCl2, 5 μg/mL insulin from bovine pancreas and 1% (v/v) penicillin/streptomycin. All cells were kept under 5% CO_2_ and at 37°C with regular mycoplasma checks. Short-term cultures (STCs) were established at NYU Langone Medical Center (Iman Osman lab) from surgically resected patient tissues (de Miera, Friedman, Greenwald, Perle, & Osman, 2012). Informed consent was obtained from all participating patients.

### 2.3 Lentivirus Production and stable cell line generation

Lentiviral particles were produced by a standard transfection protocol. 10^7^ Lenti-X™ 293T cells were plated in a 150 mm cell culture dish. Transfection was performed with psPAX2 (viral packaging plasmid) and of pCMV-VSV-G (viral envelope plasmid) and the respective cargo delivery plasmid using PEI transfecting reagent. Virus supernatants were harvested up to three days post-infection, stored at 4°C, cleared supernatants 0.2 μm filtered, concentrated using concentrating filter and stored at - 80°C until use. Target cell lines were plated each in 6 well plates using a seeding density (2.5 x10^5^ cells/well of ∼70% confluency. The cell growth medium was changed to fresh culture medium containing 8.0 μg/mL of polybrene and 50 to 75 μL of concentrated sgRNA lentivirus. 48 hours post-infection, puromycin was added to the cells in order to select stable modified cell lines (2.0 μg/mL). Stable dCas9 expression was detected by mCherry fluorescence flow-cytometry, qPCR and Western-Blot using mouse mAb (2-2.2.14) mouse anti-HA (ThermoFisher # 26183) or mouse mAb anti-Cas9 antibody (7A9-3A3, Cell Signaling). Secondary antibody was anti-Mouse IGG – HRP (A9044, Sigma Aldrich).

### 2.4 Apoptosis Assay

Expression lncRNA candidates BDNF-AS, GMDS-AS1 and XLOC_030781 was downregulated with CRISPRi alongside sgROSA non-targeting control in 501-mel-tetON dCas9 KRAB and WM1361A-tetON dCas9 KRAB as described below using best performing sgRNAs in Table S3. dCas9-KRAB cell lines were cultured in the presence of 2 μg/mL doxycycline, to induce the expression of dCas9, and subsequently 2.0 x 10^5^ cell/well were seeded in 6-well cell culture plates. On the following day, cells were infected with 50 μL sgRNA lentivirus of the respective lncRNAs in separate wells. 72 hrs post infection 150 μM of Etoposide apoptosis inducer was added to untreated cells as positive control. Cells were sgRNA infected and puromycin (2 μg/mL) selected for 48 hrs, PBS-washed and then lysed. Samples were used for immunoblotting according to (Towbin, Staehelin, & Gordon, 1979) probed with primary goat anti-PAPR antibody (Cell Signaling Technology, #9542S) and Goat Anti-Human IgG Antibody HRP conjugate (Sigma Aldrich, # AP112P). Mouse monoclonal anti-GAPDH antibody (Proteintech 1E6D9, # 60004) was applied as total protein loading control.

### 2.5 Cell Cycle Distribution Assay

The cells were treated as described in above. 72 hours post-infection cells were washed with 1x PBS and collected in FACS tubes. 3 mL ice cold 70 % ethanol were added dropwise in the cell solution while mixing and incubated for 1 hour at 4 °C. Then, cells were washed twice with 1x PBS, mixed with 200 μL staining solution containing 7-AAD (BioGem, #61410-00) according to manufacturer’s recommendation and incubated for 30 minutes at room temperature protected from light. Cells were washed twice with 1 mL 1x PBS, resuspended in 300 μL flow buffer (1X PBS/FBS 2 %, 2 mM EDTA) and subjected flow cytometry analysis (Sony, SH800S Cell Sorter or BD FACSAria). Cell cycle analysis of vital mCherry-dCas9 and GFP-sgRNA expressing double positive cells was done using FlowJo Software^TM^.

### 2.6 GFP competition Assay

Assay was done in analogy to (Wang et al., 2019) using described sgRNA RPA3 positive control and best performing sgRNAs from CRISPR screen 1.0. (Table S3). In brief, lncRNAs were knocked-down in 501-mel-tetON dCas9 KRAB or WM1361A-tetON dCas9 KRAB for up to 24 days post infection and GFP+ cell population was quantified at time points day 8, 12, 18, 20 and 24 (501mel) or day 4, 8 and 14 (WM1361a) using flow-cytometry normalized to day 4 assay start control and sgROSA. Cells were gated for vital singlets expressing mCherry-dCas9.

### 2.5 RNA In Situ Hybridization

#### 2.5.1 Preparation of fixed mammalian cells on a chambered slide

Ibidi μ-Slides 8 well slides were coated by adding 300 μL of 0.01 % poly-D-lysine for 30 minutes at room temperature. H_2_O washed slides were seeded with 2.0 x 10^4^ cells/ chamber and cells let to proliferate for 48 hours. Then chambers were washed with 300 μL of DPBS 1X. Fixation was done by adding 300 μL of 4 % formaldehyde for 10 mins at room temperature. After two additional washes with 300 μL of DPBS 1X cells were permeabilized with 300 μL of ice-cold 70% ethanol overnight at −20 °C.

#### 2.5.2 Multiplexed in situ HCR v3.0

##### Detection Stage

Ethanol was removed from the chambers and samples air dried at room temperature. The chambers were washed twice with 300 μL of 2X SSC. Samples were pre-hybridized in 300 μL of 30 % probe hybridization buffer (30 % Formamide, 5X Sodium Chloride Sodium Citrate (SSC), 9 mM Citric Acid pH 6.0, 0,1 % Tween 20, 60 μg/mL Heparin, 1X Denhardt’s Solution, 10 % Dextran Sulfate) for 30 minutes at 37 °C. Then probe solution consisting of 3 pmol of each probe in 300 μL of probe hybridization buffer (30 % Formamide, 5X SSC, 9 mM Citric Acid pH 6.0, 0,1 % Tween 20, 60 μg/mL Heparin, 1X Denhardt’s Solution, 10 % Dextran Sulfate) and incubated at 37 °C overnight. Finally, samples were washed four times 300 μL of 30 % pre-warmed probe wash buffer at 37 °C and twice for 5 minutes with 5x SSCT at RT.

##### Amplification Stage

Samples were pre-amplified in 300 μL of amplification buffer (5X SSC, 0,1 % Tween 20, 10 % Dextran Sulfate) for 30 minutes at 37 °C. Hairpin solution was prepared in two steps by separately snap cooling 18 pmol of hairpin H1 and 18 pmol of hairpin H2 (6 μL of 3 μM stock) by heating the hairpins for 90 seconds at 95 °C and then cool to room temperature in the dark for 30 min. Second step followed by adding of all snap-cooled hairpins in 300 μL of amplification buffer at room temperature. Samples were incubated for 60 minutes in the dark at room temperature. Excess of hairpins was removed by washing 5 times for 5 minutes with 300 μL of 5x SSCT at room temperature. and stained with 1 μg/mL DAPI for nuclei detection, for 1 minute and washed 3 times with 1x PBS.

##### Probe Sequences Design

Candidate probe sequences for our respective lncRNA targets were designed by Molecular Instruments ®(MI) based on their RNA sequence. Alignment to the human transcriptome was performed to minimize off-target complementarity to random genomic regions and to maximize specificity to distinctive regions of the lncRNA candidates.

### 2.6 Trans-well Migration Assay

CRISPRi 501-Mel-tetON dCas9 KRAB cell line was used to knock-down expression for 72 hrs of BDNF-AS, GMDS-AS1 and XLOC_030781 along sgROSA control using the best sgRNA each. FluoroBlok ™ 24-well inserts with 8.0 µm colored PET Membrane were coated with Matrigel at final concentration of 300 μg/mL in coating buffer solution (0.01 M Tris-HCl pH 8.0, 0.7 % NaCl). for 2 hours at 37 °C. 1.4 x 10^5^ cells were seeded per insert in 300 μL serum-free growth medium and incubated for 48 hrs. The lower chamber was filled with 700 μL complete growth medium with 1 μM Lysophosphatidic acid (LPA) chemoattractant. Invading live cells were stained with 500 μL per insert Calcein AM 2 μg/mL in HBSS 1X for 10 minutes at 37 °C before inverted fluorescent microscopy imaging. ImageJ (FIJI) software was employed to quantify migrating cells. The SAM-dCas9 501Mel cell line was generated following the previously described method (Konermann et al., 2015) by seeding 5 x 10^5^ cells per well instead.

### 2.6 quantitative PCR

qPCR was done according to (Imig et al., 2015). Typically, 1 µg of total RNA was reversed-transcribed with High-Capacity cDNA Reverse Transcription Kit (Thermo-Fisher # 4368814) and qPCR performed with GoTaq (Promega # M3001) according to instructions with primer sets listed in Table S5 and analyzed likewise *2*,^−ΔΔ*Ct*^ method (Livak & Schmittgen, 2001).

### 2.7 COMET Assay

Cell lines used for the assay were 501-mel-tetON dCas9 KRAB or WM1361A-tetON knocked-down with sgXLOC_030781_6 for 3 and 5 days post-puromycin selection start. Alkaline COMET-Assay Kit from B&D Systems (#4250-050-K) was done according to the manufacturers instruction. H_2_O_2_ treated cells were used as positive and untreated cells as negative controls. Fluorescently stained cell tails were microscopically quantified using Fiji software in percent to positive control.

### 2.8 sgRNA library construction

sgRNAs were placed in a window −300- +50 bp up- and downstream of identified transcription start site applying http://benchling.com with off target score of 70 or higher. Library targeted 262 lncRNAs upregulated in metastatic STCs vs. melanocytes and BRAFi-MEKi-resistant SK-Mel239 cells. including in total 2,761 sgRNAs (10 sgRNAs per lncRNA gene), 50 scrambled sgRNA controls and 5 lncRNA positive controls (ANRIL, BANCR, HOTAIR, MALAT1, SAMMSON). Individual sgRNA oligos were synthesized by Twist Bioscience (https://twistbioscience.com/) on a 12K array and amplified using array primers (Table S5). Using a Gibson Assembly master mix (New England Biolabs), we cloned sgRNAs into a lentiviral sgRNA GFP-tagged vector (LRG) (Addgene #65656). Gibson reactions were transformed using DH10B electrocompetent cells (Invitrogen) at 2 kV, 200Ω, and 25 µF. Bacterial colonies were quantified to obtain ∼70X coverage. Subsequently, library was deep-sequenced using MiSeq to confirm sgRNA representation. All sgRNA sequences used in this study are provided in Table S2. CRISPRa sgRNA design was done in accordance with (Konermann et al., 2015).

### 2.8 CRISPR-inhibition screen and data analysis

In brief, 501Mel and WM1361a cells were transduced with in-house generated Gibson assembled vector containing KRAB-dCas9-HA-Tag and mCherry fluorescence marker separated by a P2A cleavage signal under the doxycycline inducible TRE3G promoter in lentiviral packing signal backbone. Additionally, we cloned in opposite direction upstream of TRE3G blasticidin antibiotic marker and rtTA tetracycline resistance gene under control of the constitutive PGK promoter. Lentivirus transduced, stably selected and mCherry positive 501Mel cells used for screening (single clones D8 and D12 replicates for screen 1.0 and two cell pool replicates for screen 2.0). dCas9 expression check of 501Mel was confirmed as in Figure S2. ∼9.2 x 10^6^ cells were infected with the LRG-GFP sgRNA library expression vector at MOI 0.1-0.3 to achieve 1000x coverage. After one day of recovery infected cells were puromycin (2 µg/mL) selected for three days under simultaneous doxycycline induction (Screen 1.0) or doxycycline was added (each 2 µg/mL) after puromycin end selection after 3 days. Then puromycin was removed and followed by doxycycline addition on day 4 in order to prevent premature drop-off (Screen 2.0). Dox induction lasted for at least two days until screening start time point. One day before screening start cells were sorted near 100% positivity for mCherry and GFP double positive cells maintaining 1000x coverage before each screen as in 2.2-2.3. At time points day 7, 14 and 21 post-screen start 1000x cells of library size were collected and gDNA extracted with QIAamp DNA Blood Mini Kit (Qiagen # 511049), cleaned up with OneStep PCR Inhibitor Removal Kits (Zymo #D6030) to remove melanin as potent PCR inhibitor-Additionally, we ran in parallel non doxycycline treated cells as negative control to rule out stochastic sgRNA loss or overrepresentation. Keeping an equivalent of 1000x coverage gDNA quantity for each sample the integrated sgRNA library was PCR amplified with limiting cycles using barcoded primers listed in Table S5. Subsequent, amplified fragments were sequenced with Illumina HiSeq 2000 to yield roughly 1000 reads per guide RNA. Analysis was done similar to Shamloo et al, 2023 by cpm counting sgRNA reads of all screening time points (screen 1.0) or log2 fc read counts (screen 2.0) against control day 4 (screening start). Hit selection from screen 1.0 was done by calculating the median log2 fc of top 3 depleted sgRNAs per lncRNA at day 21 (i.e. end time point).

### 2.9 RNA-Sequencing and Analysis

RNA sequencing libraries were generated using QIAseq Stranded Total RNA Lib Kit including QIAseq FastSelect -rRNA HMR Kit and QIAseq UDI Y-Adapter Kit (Qiagen # 180743, #334385, #180310) according to the provider’s protocol. qPCR confirmed appropriate XLOC_030781 knock-down. Sequencing of the RNA libraries was performed on Illumina NovaSeq6000 PE150 with at least 30 million reads output per sample. R version 4.3.1 was used for all analyses. Raw fastq files were processed using the Zarp pipeline (Katsantoni et al., 2021) which uses FastQC, zpca and MultiQC[2(FastQC), (Ewels, Magnusson, Lundin, & Kaller, 2016) for quality control and the adapters are trimmed using Cutadapt (Martin, 2024) The reads were mapped to the human genome(hg38, Genome Reference Consortium GRCh38) using STAR (Dobin et al., 2013) and quantized Salmon (Patro, Duggal, Love, Irizarry, & Kingsford, 2017). The final output is a counts matrix which is used as input for the R package DESeq2 (Love, Huber, & Anders, 2014) to identify differentially expressed genes. Genes with a logFoldChange >1 and adjusted p-value<0.05 are used for further analysis. The volcano plot was generated using the EnhancedVolcano R-package (Blighe K, 2023) and the ComplexHeatmap (Gu, Eils, & Schlesner, 2016) package was used for generating the heatmaps. The gene sets were obtained using the msigdbr (Dolgalev, 2022) R-package and clusterProfiler (Wu et al., 2021) and enrichplot (Yu, 2023) were used for generating the GeneOntology and Gene-Set Enrichment plots.

### 2.9 ChIP-Sequencing and Analysis

ChIP-Seq was done in duplicates according to (Kloetgen et al., 2020) with modifications. In brief, 1 x 10^6^ nuclei were fixed in 1% formaldehyde in PBS and incubated at room temperature for 10 minutes, thereafter quenched by 0.125 M glycine final concentration. Cells were lysed with buffer containing 5 mM HEPES, pH 8.0, 85 mM KCl, 0.5% IGEPAL supplemented with protease inhibitors (Roche), incubation on ice for 10 minutes, and spinning to pellet the nuclei. To generate the mononucleosomal particles, the pellet was resuspended in MNase digest buffer (10 mM NaCl, 10 mM Tris, pH 7.5, 3 mM MgCl_2_, and 1 mM CaCl_2_) with 1 U of micrococcal nuclease (USB), and incubated at 37°C for 45 minutes. The reaction was stopped by adding 20 mM EDTA final and incubated on ice for 10 minutes. The nucleus was spun down and resuspended in Nucleus lysis buffer (50 mM Tris-HCl, pH 8.1, 10 mM EDTA, pH 8.0, and 1% SDS) supplemented with proteinase inhibitors (Roche). Then, the samples were sonicated using a Bioruptor (Diagenode) at high intensity for 5 cycles (30 minutes on/30 minutes off) at 4°C to obtain sheared chromatin fragments. Magnetic Protein G beads (Dynabeads) were prepared by washing them with Citrate-Phosphate Buffer (25mM citric acid, 66mM Na2HPO4, pH 5.0) and blocking them with 10 mg/mL BSA in Citrate-Phosphate buffer for 1 hour. A total of 25 µL of beads per sample and 20 µL for pre-cleaning were prepared using this method. Approximately 200 µg of chromatin fragments were pre-cleaned with 20 µL of blocked beads and 10 volumes of IP dilution buffer (167 mM NaCl, 1.1% Triton X-100, 0.01% SDS, 1.2 mM EDTA, pH 8.0, and 16.7 mM Tris-HCl, pH 8.0) for 1 hour at 4°C. 1% of the input was reserved from the chromatin. Subsequently, 25 µL of blocked beads were coupled with 5 µg of Acetyl-Histone H3 (Lys27) (D5E4) XP® Rabbit mAb or Tri-Methyl-Histone H3 (Lys4) (C42D8) Rabbit mAb (Cell Signaling) or Mono-Methyl-Histone H3 (Lys4) (D1A9) Rabbit mAb (Cell Signaling) for 4 hours at 4°C in IP dilution buffer. The antibody-coupled beads were added to the pre-cleaned chromatin in IP dilution buffer and incubated overnight at 4°C. After immunoprecipitation, DNA elution was performed by on-bead Proteinase K digest (Ambion) and overnight incubation at 65°C under high agitation. Eluted DNA was then precipitated using ethanol and glycogen. cDNA Libraries were generated as described using Kapa Hyper Prep Kit (Roche and Illumina TruSeq system) and sequenced with NextSeq500. ChIP-Seq datasets were analyzed with the HiC-bench platform (Lazaris, Kelly, Ntziachristos, Aifantis, & Tsirigos, 2017). The ChIP-Seq aligned reads were further filtered by discarding reads with low mapping quality (MAPQ < 20) and duplicated reads using picard-tools (https://github.com/broadinstitute/picard). The remaining reads were analyzed by applying the peak-calling algorithm MACS2 (version 2.0.1) 16 with input as control. Binding of histone-marks were determined from broad-peak calls.

### 2.11 Transcription Factor Enrichment Analysis

Significantly over-, and underexpressed genes (log2 fc <1, >1; adjusted p-value = 0.05) from RNA-Seq of sgXLOC_030781 knockdown at day 3 where subjected to ENCODE and ChEA consensus TFs analysis from ChIP-X (Keenan et al., 2019). TF enrichment output was plotted as -Log10(p-value).

### 2.12 ENSG00000287723 expression analysis from available datasets

Pan-Cancer expression analysis (n=5606 across 23 cancer entities), naevi and primary melanoma for ENSG00000287723 was done by retrieving fastq files from Badal et al., (2017, GSE98394), Kunz et al., (2018, GSE112509) and TCGA (GDC Data Portal access granted from Dr David Fenyo, NYU). Fastq files were trimmed using trim galore (version 0.6.6) followed by QC analysis with FastQC (version 0.12.1). Trimmed fastq were aligned to GRCh38 human genome and gene transcripts annotated with GENCODE V44 using STAR (version 2.7.7a). Normalized read counts CPM or FPKM were used to plot ENSG00000287723 expression for naevi against primary melanoma (Badal and Kunz) and primary melanoma against melanoma metastatic samples (TCGA). Statistical analysis employed unpaired t-test, two-tailed in GraphPad Prism.

### 2.13 Software and Statistics

Statistical Analysis of qRT-PCR, Western Blot quantification, Cell invasion, Luciferase and COMET assays, and finally gene expression data was performed employing a two-tailed, paired Student’s t-test to determine statistical significance: p < 0.05 (*), p < 0.01 (**), p < 0.001 (***) (GraphPad t-test calculator). Adobe Illustrator and GraphPadPrism 8.0. Fiji was utilized for microscopic object quantification related to call invasion and COMET assays. Flow cytometry analysis was done with FlowJo V8.7 TreeStar (BD Bioscience), (https://flowjo.com)

## 3. Results

### 3.1 LncRNA candidate selection for CRISPRi screen and expression of lncRNAs associated to melanomagenesis

Whole-transcriptome functional CRISPR-screening for lncRNAs is elaborate requiring large resources given the number of annotated lncRNAs reaching more than tens of thousands (Cabili et al., 2011; Derrien et al., 2012). Additionally, lncRNAs are functioning strictly context dependent and thus not all known lncRNAs are relevant in a model system of interest (Derrien et al., 2012; Djebali et al., 2012). To overcome these challenges, it appears plausible to pre-select melanoma specific lncRNAs for a functional CRISPRi focused screening based on differential gene expression (DGE). We sequenced four samples of lymph node (LN) and five brain metastasis (BM) patient derived short-term cultures (STCs) against cultured healthy donor derived melanocytes from NYU Langone Medical Center, Department of Pathology within our Melanoma Program. For convenient consecutive melanoma cell line based CRISPRi screen we added differential RNA-Seq expression analysis of 501Mel metastatic cell line (V600E) and WM1361a cell line with metastatic capacity (NRAS Q61R) to ensure ample expression of STC derived DGE lncRNAs in a stable model background. Additionally, we profiled the differential expression of lncRNAs within another clinically relevant setting namely dual BRAF- and MEK-inhibitor resistance phenotype in SK-MEL239 parental and SK-MEL239 double resistant cell line (SK-MEL239_2x, kind gift of Dr. Poulikakos. Icahn School of Medicine Mount Sinai). Overexpressed lncRNAs in SK-MEL239_2x were added to the CRISPRi library screen reasoning that genes involved in targeted therapy resistance inherit roles in melanoma cell survival and can be identified by growth (Joung et al., 2017; Konermann et al., 2015).

To ensure the relevancy of the selected lncRNAs within the experimental context, specific filters and thresholds were implemented encompassing: **a)** maintaining a false discovery rate of less than or equal to 0.05 (FDR≤+0.05 **b)** adhering to a log-fold change threshold of 0.75 or higher (logFC>+0.75) to ascertain that the lncRNAs were significantly overexpressed across clinical samples, **c)** requiring a read count per million of at least 1 (CPM≥+1) in the higher expressed cellular type, excluding genes with low expression levels which are difficult to modulate by CRISPRi, **d)** mandating that the lncRNAs carried an active histone mark, specifically H3K4me3 or H3K27ac, in the higher expressed cell type, and **e)** excluding any lncRNAs that consisted solely of mono-exonic transcripts as they might execute alternate function as for example enhancer associated RNAs (**Figure 1a**). By this we identified 88 lncRNAs to be overexpressed in BM and LN STCs vs melanocytes and 145 downregulated (**Figure 1b**), while in BM vs. LN only few lncRNAs appear to be differentially expressed (BM vs. LN 5 lncRNAs up and 12 downregulated) pointing towards similar lncRNA regulatory programs (not shown). In contrast to the STCs where only a small number of lncRNAs showed to be differentially expressed in melanoma cell SK-Mel239 vs. BRAFi/MEKi resistant derivative 174 lncRNAs are up- and 57 downregulated (**Figure S1 and Table S1**).

**Figure 1:**
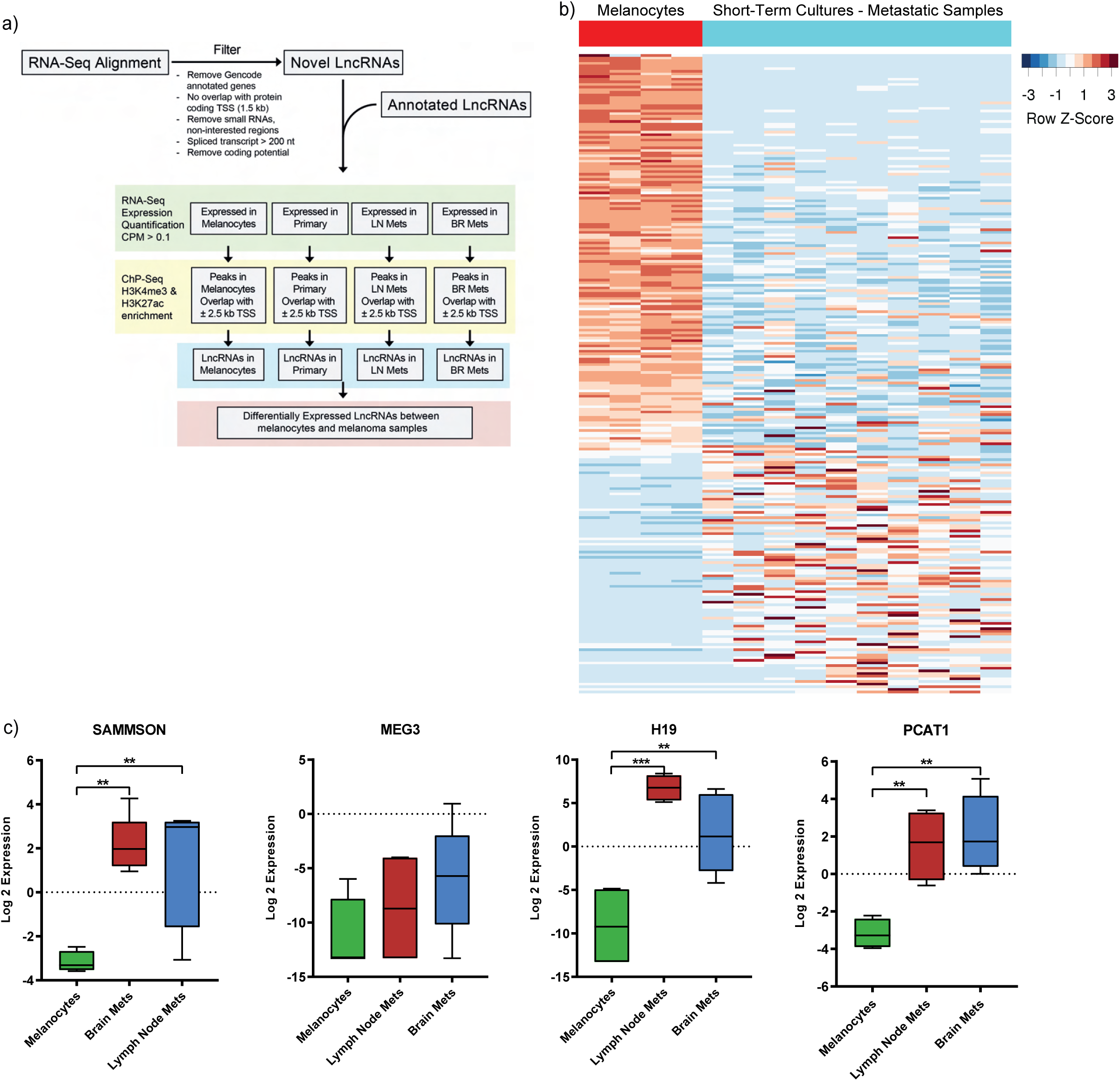
LncRNA candidate selection for CRISPRi screen and expression of lncRNAs associated to melanomagenesis. **a)** Flow-diagram of integrated RNA- and ChIP-sequencing for the pre-selection of lncRNAs. Selection parameters are FDR < 5%, logFC > 0.75 and CPM > 1 in higher expressed cell type, either Histone mark (H3K4me3 or H3K27ac) in the higher expressed cell type; no mono-exonic lncRNAs, logFC ≥ 0.5 between investigated cell type and either WM1361a or 501mel **b)** Heatmap of differentially expressed lncRNAs by RNA-sequencing in patient/donor derived melanocytes (n=4) and metastatic short-term cultures (= STC; BM = brain metastasis, n=5; LN = lymph node metastasis, n=4) and melanoma cell lines 501mel (n=4) and WM1361a (n=3). **c)** Heatmap clustering showing lncRNAs overexpressed in BM STCs vs. melanocytes and LN STCs. **d)** Heatmap clustering showing lncRNAs overexpressed in LN STCs vs. melanocytes and BM STCs. **e)** Boxplot showing the log_2_ cpm expression of selected melanoma-associated lncRNAs SAMMSON, MEG3, H19 and PCAT1 in melanocytes (n=4), BM (n=5) and LN STCs (n=4) detected by RNA-sequencing.

We further profiled our expression data for previously described lncRNAs relevant in melanoma to assure the comprehensiveness for the following CRISPRi screen. We found that except for MEG3 all analyzed lncRNAs SAMMSON, H19 and PCAT1 are at least logFC >2 overexpressed in both LN and BM STCs compared to melanocytes (**Figure 1c**). Overall, this indicates that the selected clinical samples exhibit a valid test cohort for pre-identification of additional potentially essential lncRNAs in melanoma pathogenesis.

### 3.2 CRISPR-inhibitory screen for melanoma related lncRNAs

To functionally address the overexpressed lncRNAs found in melanoma STCs we applied the CRISPR-inhibition (CRISPRi) system from (Gilbert et al., 2014; Larson et al., 2013) in a proliferation drop-off screen. We established and implemented this system in two melanoma cell line models 501mel and WM1361a derived from either metastatic lesion and primary melanoma (Hanniford et al., 2020; Marincola et al., 1994), complied a series of validation steps to assure full gene repressive performance and dCas9 expression stability. After lentiviral transduction with an adapted tetON-dCas9-KRAB-mCherry vector, selection and stable single cell line generation we tested for robust HA-tagged dCas9-KRAB protein expression in clones derived from both cell lines by Western Blot (**Figure S2a, d**). Strongest expression signal could be detected in subclones D8 and D12 in 501mel. Clones D8 and D12 were used for the first biological screening replicate 1 (CRISPRi screen 1.0) In WM1361a two clones (F2 and F11) with sufficient expression dCas9-KRAB expression were generated to be used for post-screening validations in a cell line complementary to 501mel. In line with this, strong fluorescence signal of the dCas9-KRAB co-expressed mCherry marker could be detected by flow-cytometry for all clones in both cell lines (**Figure S2b, e**). This expression signal was stable over time enabling good CRISPRi screening conditions. CRISPRi functionality in melanoma cell lines was further confirmed by an sgRNA co-expressed with GFP in a competition assay were two (501mel) or one (WM1361a) sgRNAs targeting the essential cell survival gene RPA3 (Wang et al., 2019). Loss of cellular fitness of successfully infected cells was monitored over 24 days (501mel) and 14 days (WM13161a) and compared to uninfected cell population and sgROSA negative control. The results show almost complete loss of GFP for 501mel cells while in WM1361a up to ∼85% loss of sgRNA-GFP expressing population was seen for all selected single cell clones (**Figure S2c, f**). For screen biological replicate 2.0 (CRISPRi screen 2.0) we employed a 501Mel cell pool in two independent technical replicates and cross-confirmed transduced dCas9-KRAB expressing cells by Western Blot (left), qPCR for Cas9 (middle) and mCherry fluorescence microscopy (right) of re-sorted cells (**Figure S2g**). Altogether, this indicates a fully functionally established CRISPRi system in melanoma cell lines being utilized for lncRNA loss-of-function cell viability screen.

Based on these findings, we built a customs library of 2,761 sgRNAs (10 per gene) against 88 lncRNAs up-regulated in both BM and LN vs. melanocytes and 174 lncRNAs overexpressed in BRAFi/MEKi resistant SK-MEL239 vs. parental cells. 50 non-targeting sgRNAs were incorporated as negative controls as well as positive controls against five known melanoma-associated lncRNAs (BANCR, CDKN2B-AS1, HOTAIR, MALAT1 and SAMMSON, **Table S2**). Pooled sgRNAs were subcloned into the LRG vector (Shi et al., 2015) and next-generation sequencing was carried out to confirm optimal sgRNA representation (**Figure S3a**).

Subsequently, in a 501Mel dCas9-KRAB expressing cell line we performed a loss-of-function pooled CRISPRi screen of two consecutive independent biological replicates (referred to 1.0 and 2.0). While in screen 1.0 technical replicates were comprised of 501Mel cell clones D8 and D12 (**Figure S2c**), screen 2.0 consisted of a cell pool with two technical replicates (**Figure S2g)**. Schemes are shown in **Figure 2a** (Screen 1.0) and **S3e** (Screen 2.0). After puromycin selection PCR amplified gDNA of doxycycline induced sgRNA library was collected at days 7, 14 and 21 post-screen start and subjected to deep sequencing. The end time point at day 21 post-screen start without doxycycline was analyzed as additional control to rule out stochastic library distribution changes. Thereafter, changes in sgRNA abundance were assessed by counting sgRNA reads. At first, we plotted the log2 cpm of all time points and technical replicates of screen 1.0 against the cloned sgRNA plasmid library, day 0 control and no dox control demonstrating no major sgRNA loss or random overrepresentation (**Figure S3b**). We followed up by a hierarchical clustering analysis of all Z-score normalized sgRNA counts of the same samples (**Figure S3c**). As expected we found similar sample clustering of day 0 and no dox or plasmid controls, while late time points of the screen clustered together indicating a specific response to our negative selection CRISPRi screen perturbating the expression of the selected lncRNAs. Good replicate accordance between negative control and different time point samples was achieved with R-coefficient ranging between 0.873 and 0.959 (**Figure S3d**). Principal component analysis of the screen highlighted the similarity of all control samples on PC1 (∼0.1-0.5) distinguishing them to dox induced samples over time points day 7-21 (PC1 0- -0.6), **Figure 2b**. These quality checks prompted us to select for specific lncRNA hits consistently depleted over time showcasing a potential involvement in melanoma cell survival. We achieved this by plotting the cpm sgRNA read counts of day 0 negative control over all three time points day 7, 14 and 21 of dox induced sgRNA expression (**Figure 2c**.) We implemented hit selection filter of mean fc log2>-1 of the top three depleted sgRNAs per lncRNA gene at day 21 as the time point of strongest depletion resulting in a shortlist of ten lncRNAs (**Figure S2d, Table S3)** including seven previously annotated lncRNAs and three potentially novel lncRNAs (BDNF-AS, XLOC_053861, XLOC_051428).

**Figure 2:**
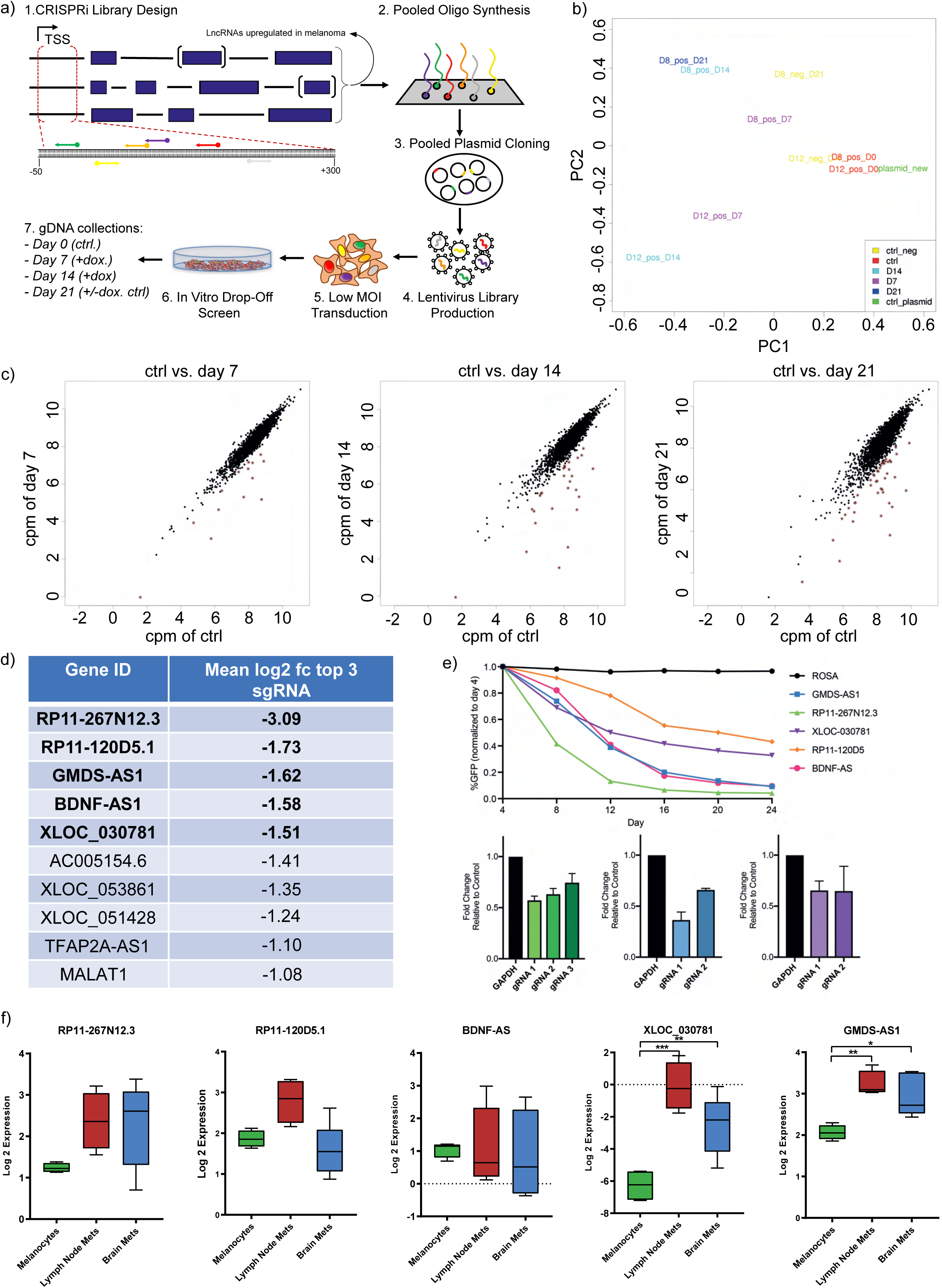
CRISPRi screen for overexpressed lncRNAs in melanoma pathogenesis and expression of selected hits. **a)** Graphical outline of the conducted CRISPRi screen in 501mel cell line. **b)** PCA plot analysis of samples used for the CRISPRi screen. D8 and D12 refers to selected 501mel-dCas9-KRAB single cell clone and D7, D14 and D21 for the days of screening time. Negative control refers to no doxycycline treatment and plasmid library for lentivirus production were added as additional control. **c)** Dot-plots shown results of CRISPRi lncRNA library screen at time points day 7, 14 and 21 post-selection and mCherry^+^/GFP^+^ cells sort as cpm of control (= day 0) vs. respective time points. Each dot represents a single sgRNA Red dots indicated significantly depleted sgRNAs and log_2_ fold-change >-1 **d)** List showing top 10 depleted lncRNA (excluding PVT1 as not validated post-screen) hits at day 21 vs. control day0 as the mean log_2_ fc of the top 3 depleted sgRNAs of each respective lncRNA **e)** CRISPRi hit validation of top 2-3 individual sgRNAs of indicated lncRNAs; top: GFP competition assay in 501mel-dCas9-KRAB from day 4 to 24 by flow-cytometry in percent GFP positive cells. sgROSA (black curve) serves as a negative control. Curves show average GFP signal loss of sgRNAs shown in bottom: bar graphs of fold change GFP expression of individual sgRNAs relative to control at day 24, color code see top panel. **f)** Boxplot showing the log_2_ cpm expression of selected lncRNA CRISPRi screening hits RP11-267N12.3, RP11-120D5.1, BDNF-AS, XLOC_030781 and GMDS-AS1 in melanocytes (n=4), BM (n=5) and LN STCs (n=4) detected by RNA-sequencing.

We validated the screen by cloning the 2-3 best screen performing single sgRNAs of the top five listed lncRNAs into LRG expression vector. We then applied a competition assay where GFP is co-expressed to the sgRNA leading to a loss of cellular fitness in case a lncRNA inhibits cell growth. We evaluated the relative GFP expressing 501mel cell population of five lncRNAs RP11-267N12.3, RP11-120D5, GMDS-AS1, BDNF-AS and XLOC_030781 over 20 days in comparison to sgROSA negative control (**Figure 2e**, top panel average depletion of used sgRNAs, bottom panel example selection of 2-3 sgRNAs CRISPRi efficiency targeting lncRNA expression relative to GAPDH). Our results confirmed that all sgRNAs are robustly depleted cells over time corroborating our screening data. We eliminated RP11-267N12.3 from our shortlist as it shares a bidirectional promoter with NUF2 mitosis related gene (DeLuca, Moree, Hickey, Kilmartin, & Salmon, 2002) making this lncRNA difficult to study and distinguish its function by CRISPRi in relation to the NUF2 neighbor gene and the previously reported oncogenic lncRNA in melanoma PVT1 was eliminated from this short list as no validation could be achieved.

To further strengthen the significance of our hit selection, we analyzed our small panel RNA-sequencing expression data set of melanocytes in LN and BM STCs (**Figure 2f**, n=4 and n=5=). We gave priority to overexpressed lncRNAs in BM and LN as a selection criterion, owing to their presumed involvement as drivers in the process of patient’s metastatic formation. Only XLOC_030781 (LN vs melanocytes: FDR = 0.000759, BM vs melanocytes: FDR = 0.00633, STC vs melanocytes: FDR = 0.00139, Table S1), GDMS-AS1 (LN vs melanocytes; FDR = 0.020, BM vs melanocytes: FDR = 0.039, Table S1) were significantly overexpressed in both BM and LN. However, RP11-120D5.1 showed only upregulation in LN. RP11-267N12. and BDNF-AS transcript in trend was also elevated LN and BM groups though not significantly overexpressed. RP11-120D5, BDNF-AS and GMDS-AS1 were additionally found overexpressed in SK-MEL239_2x, but not followed up in drug resistant context.

To further substantiate our results, we conducted a second replicate screen 2.0 with slight modifications. This time doxycycline induction was carried out later four days post-infection and two days before GFP+ and mCherry+ FACS sorting in order to minimize early drop-off of sgRNAs and therefore information loss (**Figure S3e**). Thereafter, the library processing and time points of sample acquisition was done in analogy to the first screen. The average sgRNA reads were calculated for each timepoint to the control of day 0 and the log2 fold change (fc) of the top three sgRNAs targeting each lncRNA was plotted (**Figure S3f**). Negative control sgRNAs exhibited no significant alterations in all samples (green dots), while all of our previously identified lncRNA hits were re-confirmed to be depleted at all time points. Top 50 depleted lncRNAs are highlighted in purple. Strong screening reproducibility between screen 1.0 and 2.0 was achieved as shown by the high overlap of each top 50 negatively selected lncRNA hits ranks (day 7 n=11, day 14 n=6, day 21 n=9). The most promising lncRNA candidates (BDNF-AS, GDMS-AS1, XLOC_030781) that were selected from screen 1.0 maintained comparable rankings in screen 2.0 thereby reinforcing their selection in terms of robustness and significance for melanoma cell survival (**Table S3**) and pointing to no prior drop-off due to doxycycline induction of dCas9-KRAB. Unfortunately, loss of lncRNA positive controls could only be observed in screen 2.0 for lncRNA ANRIL potentially reflecting the strong cell type dependency of most lncRNAs (Derrien et al., 2012; Djebali et al., 2012).

In conclusion, based on our screening, validation and expression analyses we carried over lncRNAs BDNF-AS, GMDS-AS1 and also XLOC_030781 to deeper phenotypic and functional characterization.

### 3.3 Phenotypic and functional characterization of lncRNA candidates

Next, the major objectives of our work were to put our selected lncRNA screening hits into relation to their drop-off behavior during the CRISPRi screen by phenotypic and functional characterization in melanoma. To do so we performed a series of in vitro cell based assays to monitor cell cycle progression, apoptosis induction, cell migration and additionally their sub-cellular localization in two melanoma cell lines to gain deeper insight into their potential mechanistic roles. Firstly, we assessed the cell cycle distribution in response to CRISPRi of lncRNAs BDNF-AS, GMDS-AS1 and also XLOC_030781 in comparison to sgROSA control by measuring the DNA content with 7-AAD dye in vital 501Mel (**Figure 3a** top left) and WM1361 cells (**Figure 3a** bottom left) using flow-cytometry of double GFP-sgRNA and mCherry-Cas9 positive cells (**Figure 3a** left). Indeed, the quantification DNA content assigned to G1, G2 and S phase (**Figures 3a** right panels and **3b**) in 501mel (top panels) indicated a substantial and significant (BDNF-AS p-value = 0.0164, GMDS-AS1 p-value = 0.0075, XLOC_030781 = 0.0477) G1 phase arrest in > 70 % of cells by knock-down of all three lncRNAs compared to sgROSA control (only ∼50% in G1 phase), which is congruent with a reduced G2 phase of about 40% in control to 14-20 % of cells under lncRNA knock-down. A similar trend for all three lncRNAs though not significant could be observed in WM1361a cell line (bottom panels).

**Figure 3:**
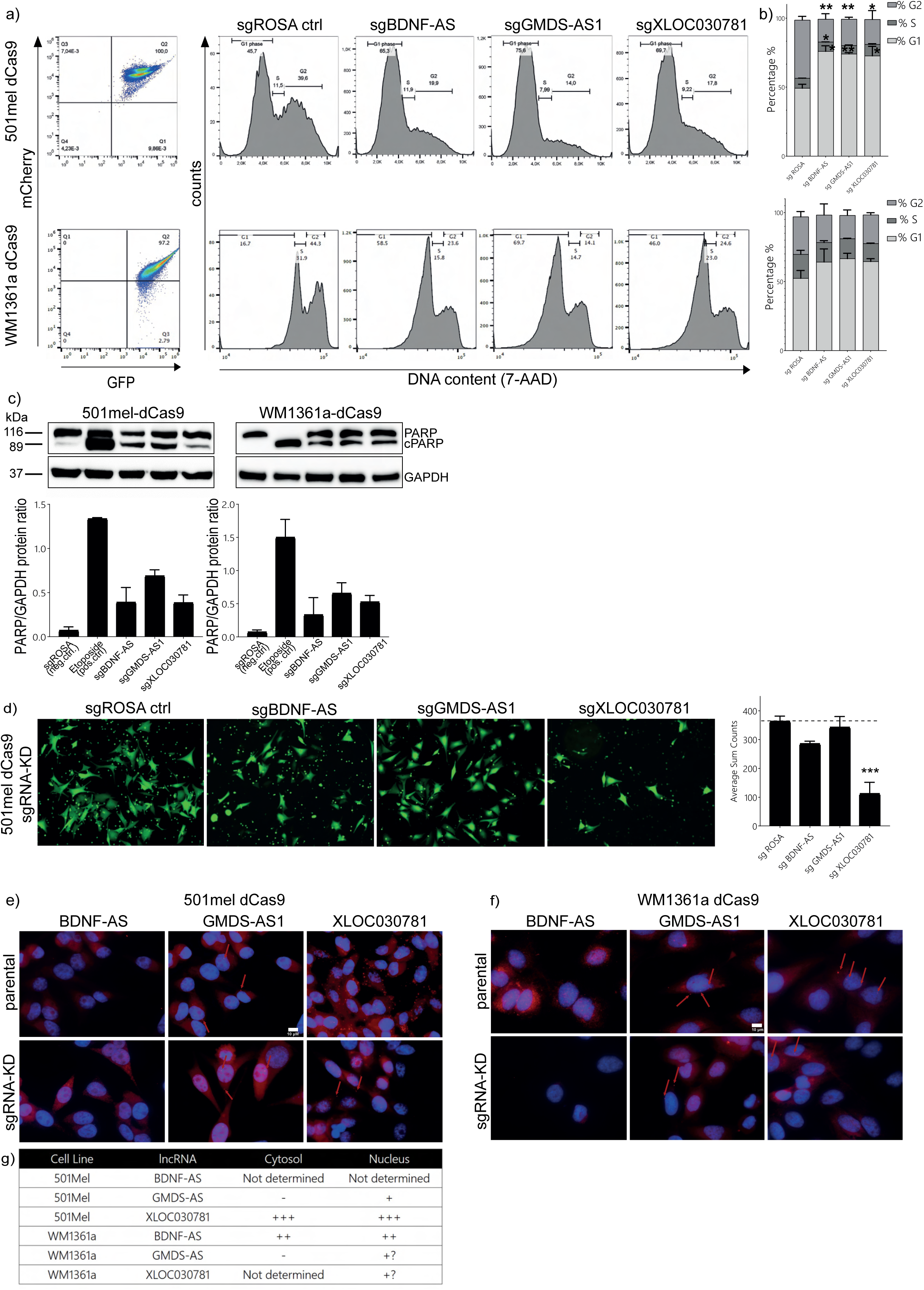
CRISPRi screen lncRNA hit functional characterization. **a)** left: flow-cytometry based dot-plot showing GFP and mCherry signal as proxy for sgRNA and dCas9 expression, right panels: 7-AAD fluorescence dependent cell cycle analysis in 501mel-dCas9-KRAB (top) and WM136a-dCas9-KRAB (bottom) cell lines upon sgBDNF-AS, sgXLOC_030781 and sgGMDS-AS1 CRISPRi knockdown. Note that top panel is plotted linear, and bottom logarithmic due to better cell cycle gating parameters **b)** Cell cycle phase quantification based on a) in percent of cells (n=2). **c)** PARP cleavage apoptosis assay performed by Western-blot upon sgBDNF-AS, sgXLOC_030781 and sgGMDS-AS1 knockdown in 501meld-Cas9-KRAB and WM1361a-dCas9-KRAB, top: representative Western-blot. GAPDH detection was used as a loading control; bottom densitometric quantification of apoptosis as cleaved PARP/GAPDH protein ratio (average of n=2). sgROSA served as negative and Etoposide apoptosis inducer as positive control. **d)** Trans-well cell migration assay. Fluorescence photomicrographs show calcein stained migrated cells of sgBDNF-AS, sgGMDS-AS1 and sgXLOC_030781 knocked-down 501mel-dCas9-KRAB cells vs. sgROSA negative control and quantification of average sum count cell numbers (n=2). **e-f)** Sub-cellular localization of lncRNAs BDNF-AS, GMDS-AS1 and XLOC_030781 in 501mel and WM1361a parental cell lines by RNA-FISH. Red arrows indicate fluorescent signals. sgRNA knockdowns serve as background staining control. Table g) summarizes sub-cellular localization and estimated signal strength.

We went on to determine the apoptosis rate upon lncRNA knock-down by examining the cleaved and uncleaved PARP protein levels by immunoblotting of mitochondrially released caspases as a hallmark of apoptosis (Boulares et al., 1999) (**Figure 3c** and **Figure S4a-d,** other replicate, densitometric quantification and qPCR knockdown confirmation). sgROSA served as negative and Etoposide apoptosis inducer as positive control in 501Mel and WM1361a. In line with the screening results and cell cycle assay we were able to detect a dramatic increase of cleaved PARP in both cell lines for all three lncRNAs BDNF-AS, GMDS-AS1 and also XLOC_030781, illustrating the importance of their expression in cell survival.

Ultimately melanoma lethality predominantly arises from the occurrence of metastasis, wherein the extensive cell dissemination plays a major role (Schadendorf et al., 2015). As a simple in vitro approximation, a trans-well migration assay is considered to reflect the migration and invasion of cells., We performed this assay under lncRNA knockdown condition for BDNF-AS, GMDS-AS1 and also XLOC_030781 using the best performing sgRNAs in 501mel cells. Interestingly, only knock-down of XLOC_030781 could attain a robust reduction of trans-well migrating cells by ca. 60 % (**Figure 3d**) eventually in line with its high overexpression found in BM and LN STCs **(Figure 2f**). As no difference in migrative phenotype was seen for GMDS-AS1 and BDNF-AS repression we performed only for XLOC_030781 another trans-well assay replicate. Likewise, a strong reduction in cell mobility was confirmed for sgXLOC_030781 compared to control (**Figures S4e, f**). Further on, we tried to reverse this anti-migrative phenotype by CRISPR-activating BDNF-AS and XLOC_030781 in 501mel. Even though strong expression upregulation for both lncRNAs with each three different sgRNAs ranging between several ten- to up to hundred-fold was accomplished, no obvious difference in the trans-well assay was observed which is possibly due to already saturated expression levels in cells (**Figures S3g-i**).

An essential step in functional understanding of lncRNAs is to determine their sub-cellular localization. We therefore carried out multiplexed RNA In-situ hybridization using fluorescently labelled anti-lncRNA probes and analyzed their localization by microscopy (**Figure 3e, f**). The CRISPRi cell lines 501mel-tetON dCas9 KRAB and WM316A-tetON dCas9 KRAB knocked-down for BDNF-AS, GMDS-AS1 and also XLOC_030781 were used as background controls, while the parental cell lines 501mel and WM1361a were stained to determine the lncRNA localization. **Figure 3g** table summarizes the localization of each lncRNA in both cell lines. In 501mel strong signal for XLOC_030781 could be detected in both nucleus and cytoplasm compartment, while GMDS-AS1 was present only in the nucleus and BDNF-AS could be not detected in neither nucleus nor cytoplasm. In WM1361a a weak signal for XLOC_030781 was found in nucleus only comparable to the pattern seen for GMDS-AS1, while BDNF-AS expression was robustly detected in the nucleus and cytoplasm. This reflects the generally lower expression of those lncRNAs in WM1361a relative to 501mel (**Table S1**). Cell staining of knocked-down lncRNAs were strongly reduced in signal intensity showcasing the specificity of RNA-ISH approach.

I sum, we phenotypically characterized three CRISPRi lncRNA screening hits and their importance in cell survival and invasive potential in vitro.

### 3.4 XLOC_030781 CRISPRi transcriptome profiling

After having established that our CRISPRi screen resulted in valid lncRNA hits contributing to cell growth, cell cycle apoptosis and migration in melanoma we decided to deeper classify the molecular consequences on XLOC_030781 expression at the transcriptomic level. We selected XLOC_030781 for several reasons: **1. Figure 2f** shows upregulation both in BM and LN STCs melanoma reflecting unique features eventually relevant for site independent metastatic lesion formation **2.** We verify XLOC_030781 lncRNA contributing to cell growth (**Figures 2e, 3a, b**) and survival (**Figure 3c**) and **3**. we established an invasive-like phenotype in melanoma (**Figure 3g**) outstanding the two other lncRNAs tested. **4.** at the time of study XLOC_030781 was a novel lncRNA and its dual localization in the nucleus and cytoplasm (**Figure 3e**) suggests a compartment shuttling or multi-functional roles, which is interesting to study.

To perturbate XLOC_030781 lncRNA expression the most effective sgRNAs (sgRNA6 and sgRNA7) were employed each in technical duplicates including two negative control samples in 501-mel-tetON dCas9 KRAB cells were dox treated for 3 and 5 days day. Ribo-depleted RNA converted cDNA was then subjected to deep-sequencing and differential transcriptome analysis. PC analysis showed well separation of all sgXLOC_030781 treated against control samples at both time points on PC1 explaining 67% variance (**Figure 4a**). Differential gene expression analysis showed a dramatic and significant transcriptome divergence upon lncRNA knockdown compared to control at both time points (**Figure 4b**, and **Figure S5a** log2 fc = 1, p-value 0.05) while there were only marginal expression differences between sgRNA knock-downs at day 3 and 5 (**Figure S5b**). In total we observed 1294 genes up- and 1462 downregulated at day 3 and 986 up- and 1179 down-regulated at day 5. Subsequent Gene ontology (GO) term enrichment analysis of differentially expressed genes (**Figure 4c**, top 20 GO terms up- and down genes) at day 3 highlighted features such as DNA replication-dependent nucleosome assembly and DNA replication-dependent nucleosome organization in the downregulated genes. This is might implicate XLOC_030781 functioning on cell division and proliferation at the transcriptional level. In the up-regulated genes GO terms like ER to Golgi vesicle mediated transport or autophagosome organization appeared (also shown in GSEA plots in **Figure 4d**). Of note, GO terms of up-regulated genes were quite disperse to the down-regulated gene set consisting the first ones mainly of cell functions located in the cytoplasm while latter one’s functions take place predominantly in the nucleus. This could eventually be attributed with the presence and functionality of this lncRNA in both compartments (**Figure 3d-f**). In general, the same pathways and gene set enrichment analyses at day 5 resembled well the ones day 3 and are summarized in **Figures S5c, d** confirming our data.

**Figure 4:**
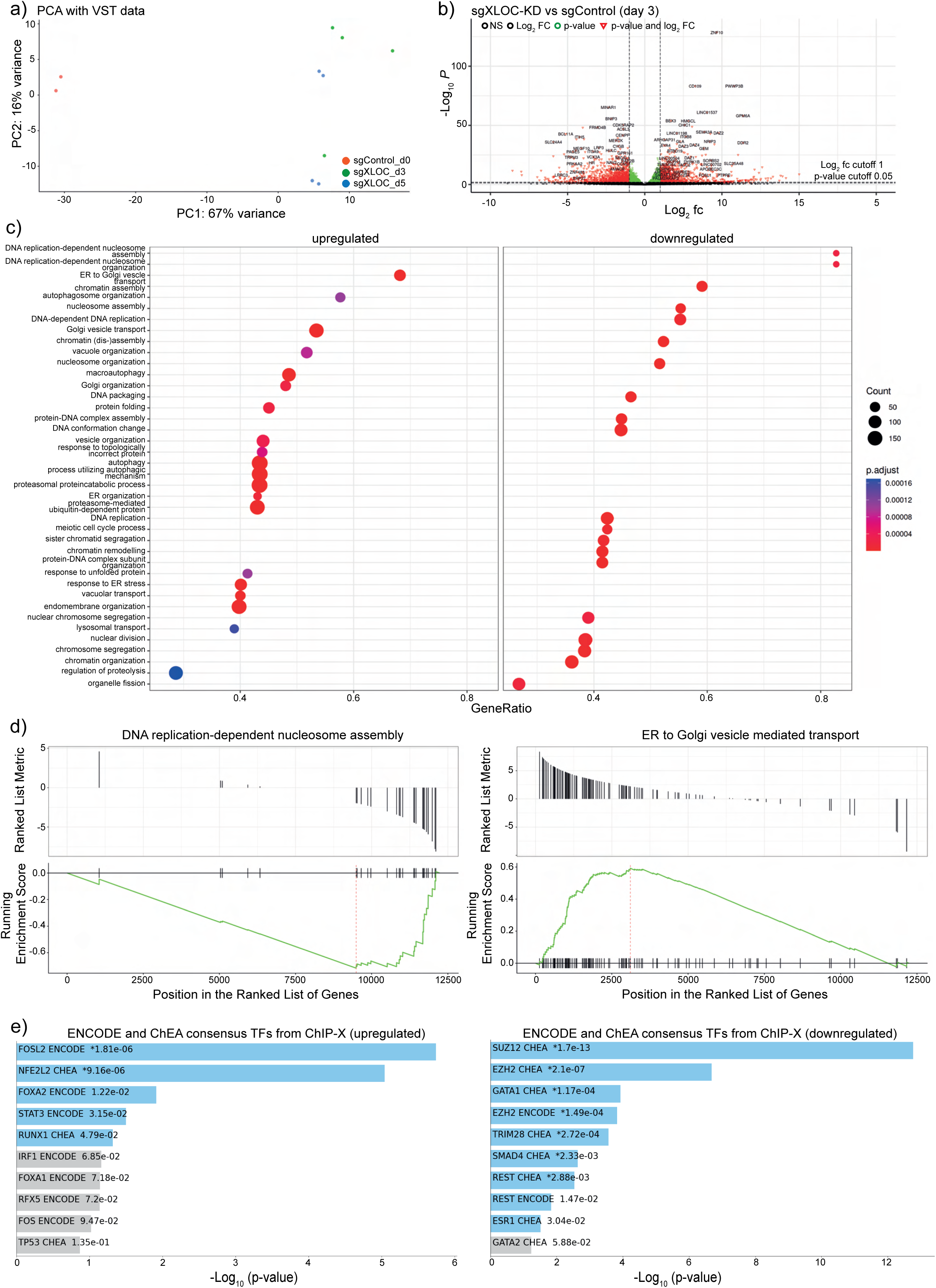
XLOC_030781 CRISPRi knockdown and RNA-sequencing transcriptomic expression analysis in 501mel-dCas9-KRAB. **a)** PCA analysis of sgXLOC_030781 CRISPRi at day 3 and 5 post-infection vs. day 0 negative control (n=2). **b)** Volcano plot of significantly differentially expressed genes (red) upon sgXLOC_030781 knockdown at day 3, p-value cut-off 0.05, log_2_ fc = 1. **c)** Gene ontology enrichment analysis of up- and down regulated gene-sets upon sgXLOC_030781 knockdown at day 3. **d)** GSEA enrichment analysis of each top example of up-regulated (ER to Golgi vesicle transport, left) and down-regulated genes (DNA-replication dependent nucleosome assembly, right). **e)** Up-stream transcription factor analysis of up-and down-regulated genes using ChEA upon sgXLOC_030781 knockdown at day 3.

We next aimed to delineate potential molecular functions of XLOC_030781 from gene signatures obtained by our knock-down RNA-sequencing approach. Recently, it has been proposed that certain lncRNAs functionally act to cis-regulate their neighbor gene expression by shaping the proximal chromatin environment (Gil & Ulitsky, 2020). To check this hypothesis, we collected the expressed 14 neighbor genes out of a window approximately one mega-base up- or downstream of XLOC_030781 locus and plotted the z-score normalized gene expression of sgRNA treated vs. control samples at all time points. We found no significant expression correlation between those comparisons ruling out the possibility that XLOC_030781 serves as a cis-regulator at the neighboring chromatin (**Figure S5e).** We then meta-analyzed the enrichment of differentially expressed genes for their occupancy and regulation by different up-stream transcription factors using ENCODE and ChEA consensus TFs from ChIP-X a transcription factor target library relying on deposited ChIP-Seq data (Keenan et al., 2019). Top two p-value ranked enriched transcription factors among the XLOC_030781 knock-downed genes included FOSL2 and NFE2L2 transcription factors and among the upregulated genes SUZ12 and EZH2, both components of the PRC2 repressor complex (Sauvageau & Sauvageau, 2010). Although all these transcription factors are already associated also as drivers of melanoma (Chen et al., 2021; James et al., 2023; Jessen et al., 2020; Terranova et al., 2021; Verma et al., 2022) their exact implications in XLOC_030781 phenotype drivers remain inconclusive.

Since an extraordinary high number of genes were misregulated upon XLOC_030781 knock-down we asked the question whether this might be an effect due to impaired DNA damage repair or reduced genome stability. The COMET assay is a single cell approach to quantify affected DNA damage repair and DNA fragmentation accumulation. We performed this assay upon sgXLOC_030781 condition at two different time points day 3 and 5 post sgRNA infection in 501mel and WM1361a cells in duplicates. Non-treated cells served as negative and DNA oxidizing agent H_2_O_2_ treated cells as positive control.

Tail generating cells were quantified by microscopy and in 501mel treated with sgRNA showed a significant higher proportion of cells with a COMET (>20%, p-value = 0.0076, student’s t-test, two-paired) at day 3 as compared to control (ca. 5%). All the other time points and cell lines showed a similar trend so not significant could be observed **(Figures S5 f, g**). This indicates that XLOC_030781 might be involved in genome stability DNA and damage repair.

### 3.5 TCGA data mining of XLOC_030781/ENSG00000287723

During the course of the manuscript preparation it turned out that lncRNA XLOC_030781 became annotated as ENSG00000287723. We therefore were able to mine clinically relevant TCGA expression data sets for XLOC_030781 and clinical associations thereof. First, we profiled XLOC_030781 expression in 5606 cancer samples over 23 tumor entities and found this lncRNA to be very skin cutaneous melanoma (SKCM) specific (**Figure 5a**). Next, we turned our attention to two published data sets from (Badal et al., 2017) and (Kunz et al., 2018) in order to stratify XLOC_030781 expression in primary melanoma (n=51 Badal et al. and n=57 Kunz et al.) and naevi biopsies (n= 27 Badal et al. and n=23 Kunz et al.) which is in line with our small scale in-house expression analysis (**Figures 2f and 5b**). In contrast to our initial findings, TCGA data analysis of SKCM metastatic lesions (n= 367) in comparison to 76 primary melanomas didn’t identify a significant expression difference of XLOC_030781 between the two patient groups (**Figures 5c and 2f**). Additionally, high expression of XLOC_030781 appeared to be associated with a significantly higher patient survival probability (p-value = 0.0216) as compared to lowly expressing melanoma patients (**Figure 5d**). This might be attributed to the pleiotropic and strong context dependent functions generally found among lncRNAs (Derrien et al., 2012; Djebali et al., 2012).

**Figure 5:**
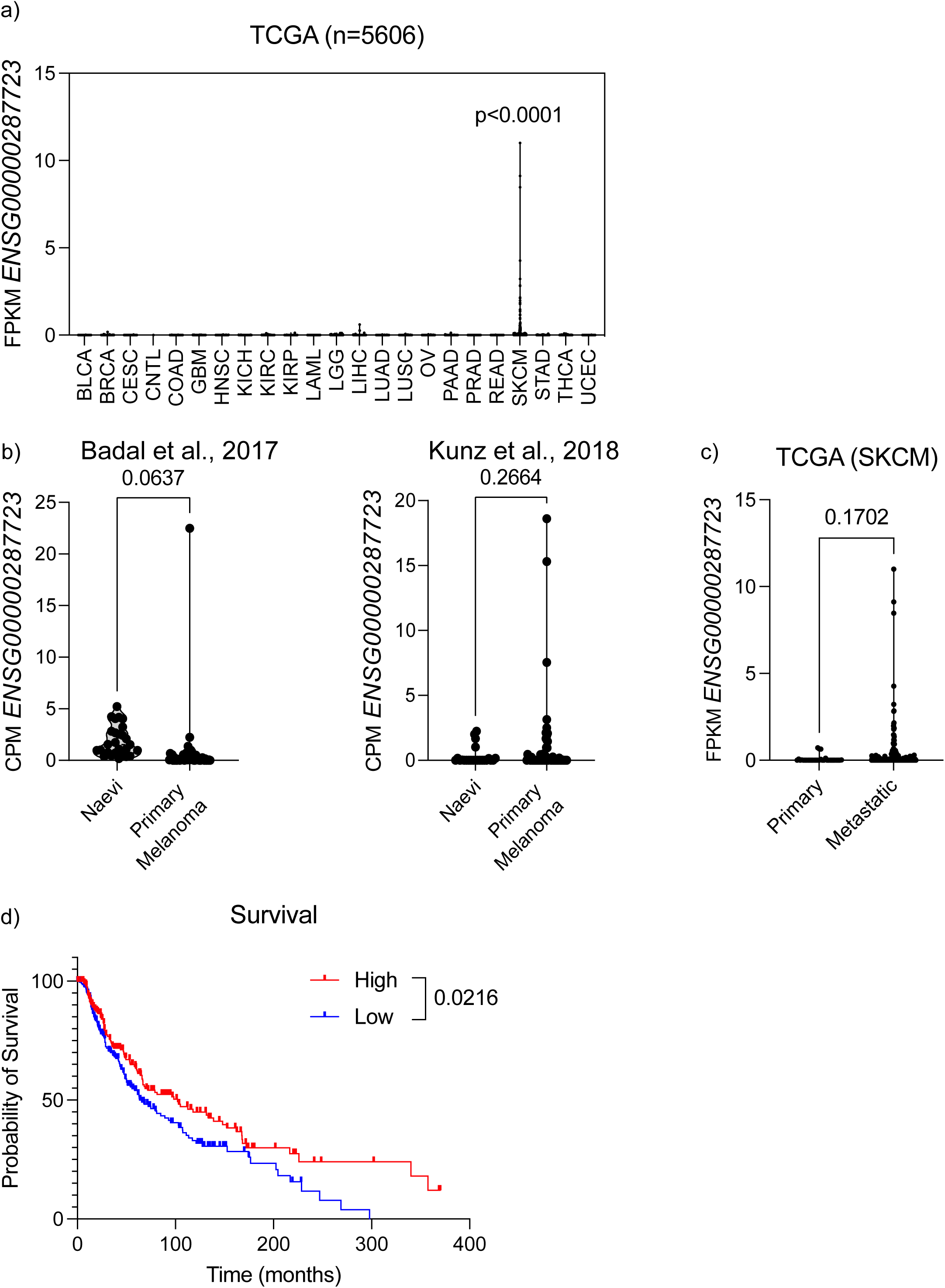
TCGA XLOC_030781 expression data analysis in cancer and clinical melanoma samples **a)** Pan-cancer expression analysis using TCGA. **b)** XLOC_030781 expression analysis in RNA-Sequencing data analysis from Badal et al., 2017 naevi (n=27) vs. primary melanoma (n=51) (left), Kunz et al., naevi (n=23) primary melanoma (n=57)(right), unpaired t-test and **c)** TCGA of primary melanoma (n=76) vs. metastatic melanoma (n=367) **d)** Kaplan-Meier patient survival in XLOC_030781 high vs. low expression samples (Log-rank Mantel-Cox test).

Next, to better understand biological features of patients carrying high and low expression of XLOC_030781 (**Figure S6a**), we performed DeSeq2 analysis on a subset of TCGA melanoma patients (High=20, Low=125) and found 3656 and 4458 genes significantly associated to high (XLOC^High^) and low (XLOC^Low^) expression of XLOC_030781 respectively (adjpval<0.05) (**Figure S6b,c**). Gene ontology analysis of these genes shown a significant association with neural-like, angiogenesis and calcium channel genes in XLOC^High^ patients. On the contrary, XLOC^Low^ patients do express genes involved in keratinocyte biology and pigmentation (i.e., MITF, TYR, TYRP1) (**Table S7**).

## 4. Discussion

The current study examines the expression of lncRNAs in metastatic melanoma derivatives compared to healthy cells, as well as in various melanoma cell lines. The goal is to lay the foundation for a focused CRISPR inhibition screening of lncRNAs involved in proliferation and invasion. Our hypothesis was that some of these lncRNAs may have oncogenic gain-of-function properties, potentially increasing the chances of identifying new ones associated with metastatic melanoma while minimizing screening efforts. Through this approach, we identified thousands of expressed lncRNAs, including many novel ones. Among these, 262 were significantly overexpressed, significantly expanding the known repertoire of lncRNAs associated with metastatic melanoma (Melixetian, Pelicci, & Lanfrancone, 2022), potentially holding relevance for disease pathology. Furthermore, we successfully validated the phenotypic effects of four of the top ten hits from the screening (GMDS-AS1, BDNF-AS, RP11-267N12.3, and XLOC_030781). Three of these were further characterized, demonstrating their positive impact on cell cycle regulation, apoptosis, and invasion in vitro. Our primary candidate, XLOC_030781, underwent additional analysis, revealing significant transcriptional changes upon perturbation. This suggests its involvement in bi- or multicellular functions related to nucleosome assembly and vesicle transport.

In our initial CRISPRi screen 1.0, we identified 12 potential lncRNA hits from the 262 lncRNAs present in the library (see Table S3). These hits were largely confirmed by a robust 40% overlap of the top 50 hits in our second screening replicate, which exhibited more stringent performance (Figure S3g). This reinforces the evidence for the involvement of these top lncRNAs in crucial molecular pathways supporting cell survival or proliferation. Overall, this resulted in a raw hit rate of 4.5%, comparable to our previous work on lncRNAs in melanoma BRAFi-resistance using a similar pre-selection strategy (Shamloo et al., 2023). This underscores the potential of our approach, which we believe can be applied to other cancer types and screening strategies. However, hit validation was successful for only 5 out of 6 lncRNAs in our GFP competition assays, leaving some false positive hits. Additionally, two more validated lncRNAs were excluded from functional studies due to low performance or biological obstacles (RP11-120D5 and RP11-267-N12.3). Nevertheless, our screening effort vastly reduces compared to existing whole-genome off-the-shelf lncRNA libraries like CRINCL (Liu et al., 2017), which consist of several specialized sub-libraries or a few general libraries, each containing over 1000 genes and >10,000 guides. The absence of selected lncRNA positive controls within the range of hit depletion, other than ANRIL in screen 2.0 (Figure S3f), can be attributed to two factors: either their downregulation does not affect cell survival, or compensatory mechanisms involving alternative genes ensure cellular viability in our model cell lines. This highlights the careful consideration and selection of phenotypically active ncRNAs for future CRISPRi screens, or their inherent versatility and transferability per se.

Our functional characterization studies prioritized three lncRNAs (BDNF-AS, GMDS-AS1, XLOC_030781) that ranked highly (among the top 10) in both CRISPRi screens, based on their performance in the validation assay and their expression pattern in our in-house small-scale melanoma cohort (Figure 2e and f). BDNF-AS (Brain-Derived Neurotrophic Factor Antisense Strand RNA) is located on chromosome 11 (genomic coordinates from GRCh38.p14 assembly: chromosome 11: 27,506,830-27,698,231 forward strand) and has been implicated in various cancer types. Interestingly, while it’s consistently downregulated across cancers like colorectal cancer, esophageal cancer, cervical cancer, glioblastoma, and osteosarcoma (Ghafouri-Fard, Khoshbakht, Taheri, & Ghanbari, 2021), its role in melanoma remains unexplored. On the other hand, GMDS-AS1 (GDP-mannose 4,6-dehydratase Antisense 1) is a novel lncRNA transcribed from a genomic locus on chromosome 6 (genomic coordinates from GRCh38.p14 assembly: chromosome 6: 2,245,718-2,525,976 forward strand). To date, studies have shown its antitumorigenic properties in lung adenocarcinoma and hepatocellular carcinoma, while promoting tumorigenesis in colorectal cancer (Huang, Zhong, Li, Wang, & Qin, 2023; Peng et al., 2023; Ye et al., 2023). Similar to BDNF-AS, its association with melanoma has not been explored until now. Our findings suggest that both BDNF-AS and GMDS-AS1 tend to be overexpressed in metastatic melanoma, affecting cell cycle progression and apoptosis but not migration. This indicates a role in cell survival rather than metastasis. However, further analysis of expression data from larger patient samples is needed to confirm our observations beyond our presented dataset (see Figure 2f).

Lastly, XLOC_030781 represents an unannotated lncRNA located on chromosome 18 (genomic coordinates from GRCh38.p14 assembly: chromosome 18: 15,164,633-15,164,933). Designated as ENSG00000287723 since September 2023, this lncRNA remains largely unexplored with existing information is limited to its expression in various cell types and tissues, including the cortical plate, ventricular zone, ganglionic eminence, and 68 other cell types or tissues (“AP005901.6 expression in human,”) However, among the 23 analyzed cancer types, XLOC_030781 seems to exhibit the typical lncRNA expression pattern specific to melanoma, suggesting a particular function in this context. While small-scale lncRNA expression assessment using metastatic STCs (Figure 1b and 2f) remains crucial, there are conflicting indications in larger melanoma cohorts of different stages (Figure 5c). This doesn’t necessarily reflect the invasive or migratory capacity observed at least in vitro for lncRNAs BDNF-AS and GMDS-AS1, but it does for XLOC_030781. However, further clarification is needed regarding their in vivo metastatic phenotype. Nevertheless, XLOC_030781 expression appears to be tightly associated with migratory capacities, at least in a subset of patient samples. Interestingly, melanoma patients with high expression levels of XLOC_030781 showed a significantly better survival probability compared to those with low expression levels. This phenomenon might be attributed to the potential multi-functional roles of this lncRNA, where its invasive phenotype is balanced by immunoregulatory effects, as observed in pathway hallmark analysis of patient data expression (Table S7). We observed significant upregulation of gene sets involved in ER to golgi vesicle transport upon repression of XLOC_030781 (Figure 4c). This process is critical and could potentially influence HLA surface expression and recognition by the host immune system (Pishesha, Harmand, & Ploegh, 2022).

However, we clearly delineated the oncogenic features of XLOC_030781 in our in vitro test systems, demonstrating a positive impact on cell proliferation, migration, and cell cycle progression, while negatively affecting apoptosis (Figure 3). Yet, uncovering the exact mechanisms and determining whether these phenotypes are relevant in vivo or in situ remains unclear, necessitating further studies in pre-clinical models for clarification. The multifunctional roles of lncRNA XLOC_030781 in melanoma are evident for several reasons: it is present in both the nucleus and cytoplasm in roughly equal proportions, and discrete gene set enrichments of transcripts upon knockdown are observed. Upregulated transcripts are mainly cytoplasmic-based, while downregulated transcripts are apparent in the nucleus. Moreover, the substantial number of differentially expressed genes in our RNA-sequencing analysis of XLOC_030781 raises intriguing questions about its underlying principle. One possibility is that XLOC_030781 orchestrates the function of critical master regulators in transcription or acts on epigenetic regulation. However, our conducted transcription factor analysis (see Figure 4e) did not provide a clear answer to this. Alternatively, XLOC_030781 may safeguard genomic integrity, as suggested by GSEA of downregulated transcripts related to DNA-dependent replication or nucleosome assembly (Figure 4c and S5c), along with our COMET assay findings (Figure S5f) may point towards the latter direction (Branzei & Foiani, 2010; De Marco Zompit & Stucki, 2021; Saxena & Zou, 2022). However, the increased number of COMETS may merely reflect a late effect on apoptosis directed by XLOC_030781. In general, disentangling the function of XLOC_030781 in metastatic spread versus cell survival while retaining its potential positive immunogenic function could be a valuable strategy for targeting this lncRNA. However, extensive genomic, biophysical, and in vivo work is required in the future to holistically understand the multifunctional roles of XLOC_030781 and its implications for melanoma progression and pathogenesis.

In conclusion, we provide a powerful set of potentially relevant and novel lncRNAs in melanoma, while the usefulness of certain of the identified lncRNAs as biomarkers in disease or even future therapeutic target structures remains to be determined.

## Acknowledgments

We acknowledge A. Heguy and the NYU Genome Technology Center for expertise with sequencing experiments; the NYU Flow Cytometry facility for expert cell sorting; the Applied Bioinformatics Laboratory for computational assistance. We thank all past and present members of the Aifantis, Hernando and Imig Labs for excellent scientific discussion, continuous support and intellectual contributions. We are grateful to Prof. Bastiaens MPI Dortmund and Prof. Daniel Summer and Tzu-Chen Lin, TU Dortmund granting access and help to their fluorescent microscopy and flow-cytometry infrastructure. We appreciate the technical support and reagents supply for the RNA-FISH by Dr. C. Schröter MPI Dortmund. We thank Prof. P. Poulikakos Icahn School of Medicine at Mount Sinai for supply of SK-MEL239_2x double resistant cell line. This work has used computing resources at the High-Performance Computing Facility at the NYU Medical Center and at “Gesellschaft für wissenschaftliche Datenverarbeitung mbH Göttingen” (GWDG).

## Author’s Contributions

J.I., I.A., and A.T. conceived, designed and supervised the study. SS and S.T. conducted bioinformatics analysis. J.I., K.H., E.W., P.T., E.S.B., E.V.S.M. and S.P. executed experiments. I.O. provided essential pathological material. P.B. meta-analyzed TCGA data. T.Z.L and M.S. performed FACS and cell sorting experiments with S.P.

## Supporting Information

Supplemental Results and Methods

Table S1: DEG_genes_STCvsSamples_RNA-ChIP-Seq

Table S2: sgRNA_CRISPRi_screen

Table S3: CRISPRi_screens

Table S4: XLOC303781_KD_RNA-Seq

Table S5: Oligonucleotides

Table S6: ENSG00000287723_Badal_Kunz

Table S7: Deseq2_ENSG00000287723_TCGA

### Data Availability and Resources

#### RNA-Sequencing

RNA-Seq of melanoma short-term cultures and cultured melanocytes: Hanniford et al., Cancer Cell, 2015 (GSE138711) and Fontanals-Cirera et al., 2017 (GSE94488).

RNA-Sequencing of melanoma cell lines: WM1361a (WM1361a-pWPI) and SK-MEL239: Hanniford et al., Cancer Cell, 2015 (GSE138711), 501mel (501mel-JQ1R-24hr, 501mel-JQ1R-6hr) Fontanals-Cirera et al., 2017 (GSE94488). SK-MEL239_2x (BRAFi and MEKi resistant) this publication: https://doi.org/10.17617/3.5S92GO

501mel XLOC303781_KD_RNA-Seq: https://doi.org/10.17617/3.5S92GO

#### ChIP-Sequencing histone modifications

ChIP-seq of melanoma STCs: Verfaillie et al., 2015 (GSE60666),

ChIP-seq of cultured melanocytes Fontanals-Cirera et al., 2017 (GSE94488)

ChIP-seq of melanoma cell lines 501mel (Fontanals-Cirera et al., 2017, GSE94488), WM1361a, SK-MEL239 and SK-MEL239_2x: this publication: https://doi.org/10.17617/3.5S92GO

#### CRISPR library screen sequencing

CRISPRi screen 1.0 and 2.0: this publication: https://doi.org/10.17617/3.5S92GO https://doi.org/10.17617/3.5S92GO can be found at https://edmond.mpg.de/dataverse/edmond a public large data repository provided by the Max Planck Society.

## Supplemental Figures

**Figure S1:**
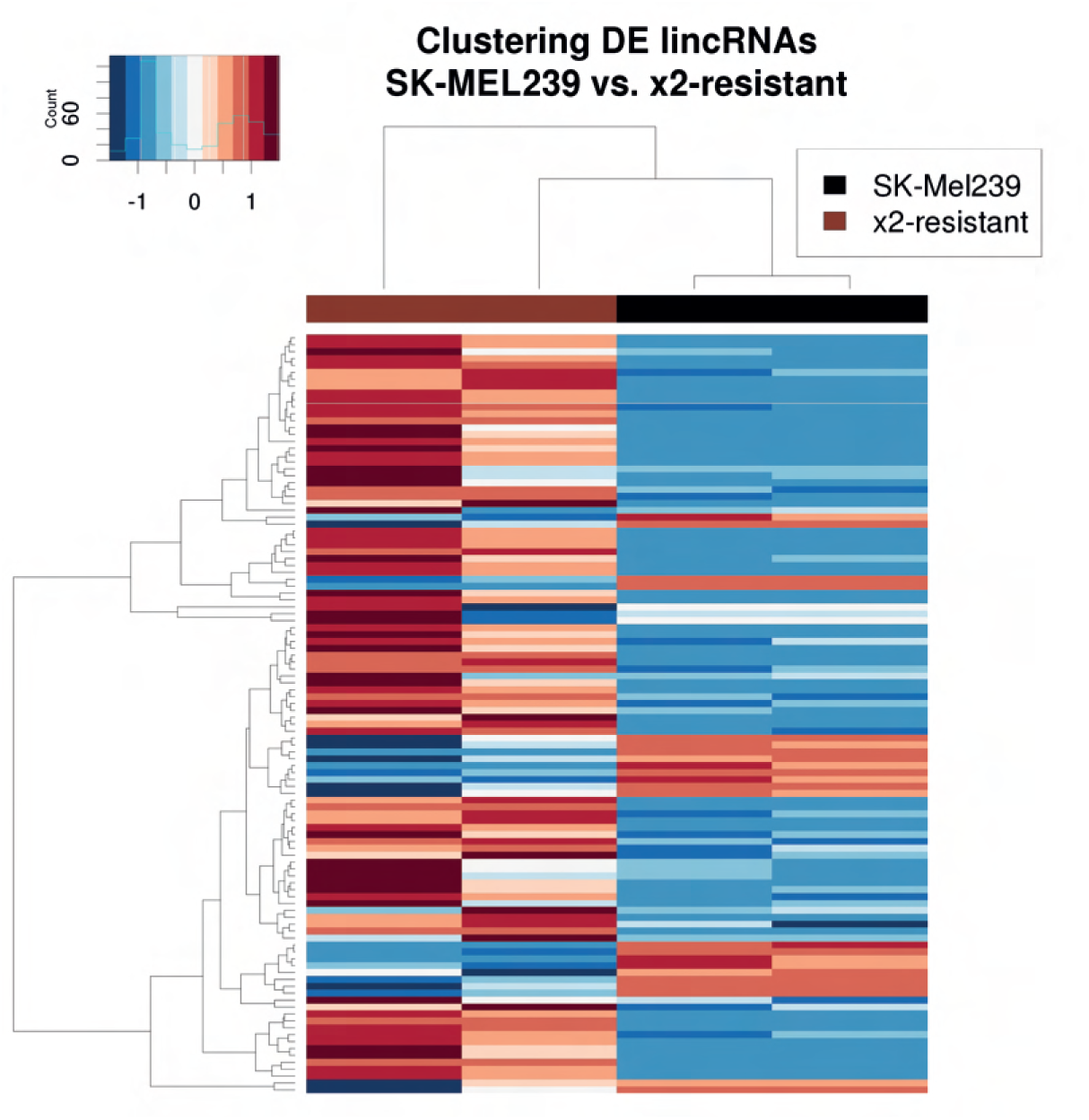
Heatmap showing differential lncRNA expression in SK-Mel239 melanoma cell line vs. BRAF-and MEK-inhibition resistant cell line (n=2).

**Figure S2:**
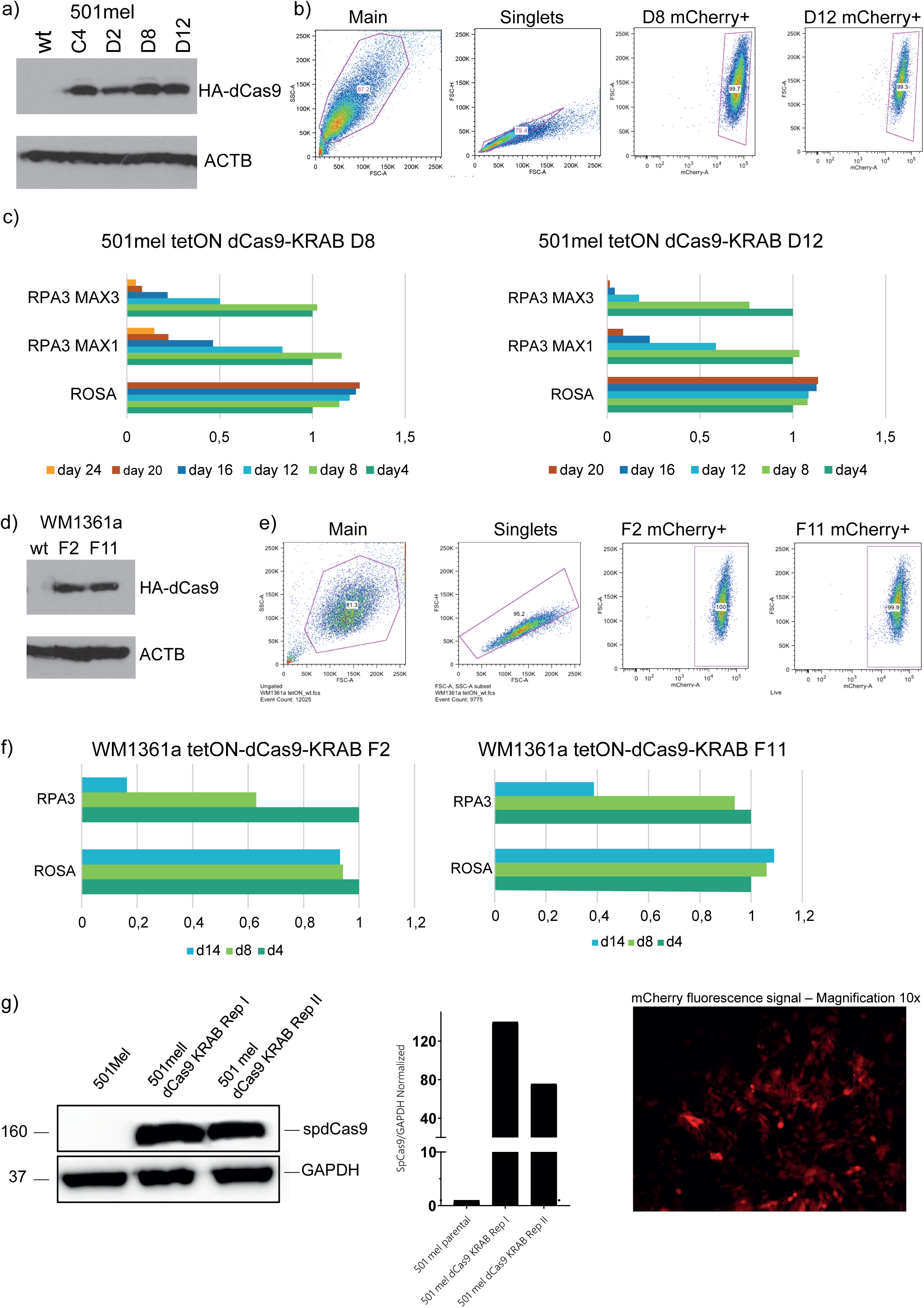
CRISPRi system establishment in melanoma cell lines. HA-tagged dCas9-KRAB protein expression in diverse single cell clones by Western-blot, mCherry-dCas9-KRAB by flow-cytometry and GFP competition assays for 2 sgRNAs targeting essential gene RPA1 and sgROSA negative control over indicated time day 4 to day 20 in **a)- c)** 501mel single clones D8 and D12 and **d)-f)** WM1361a F2 and F11 over day 4 to day 14. **g)** dCas9-KRAB expression confirmation in 501mel cell pool by Western-blot and mCherry fluorescence detection for repetition of CRISPRi screen 2.0. (related to supplemental Figure S3).

**Figure S3:**
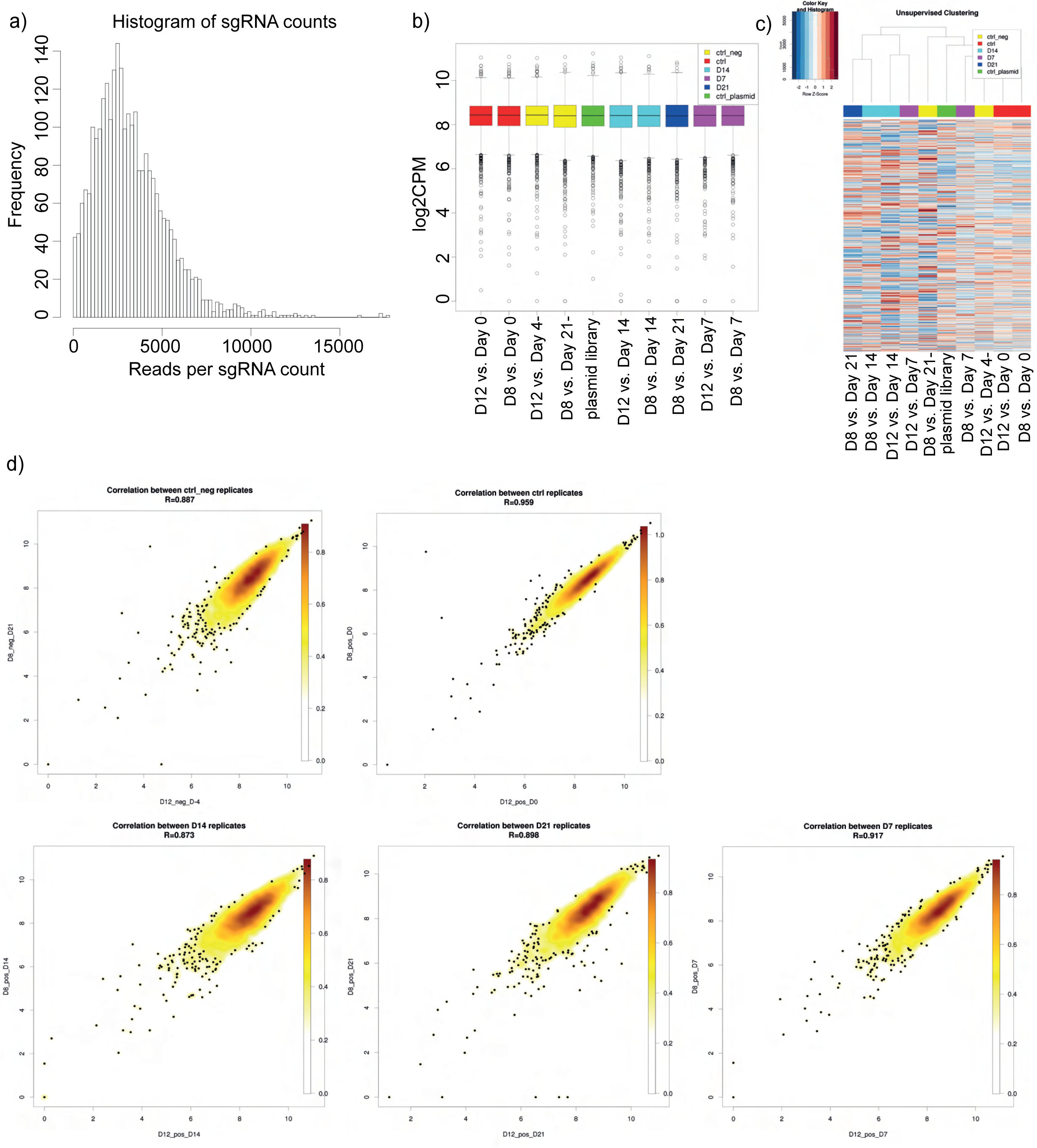
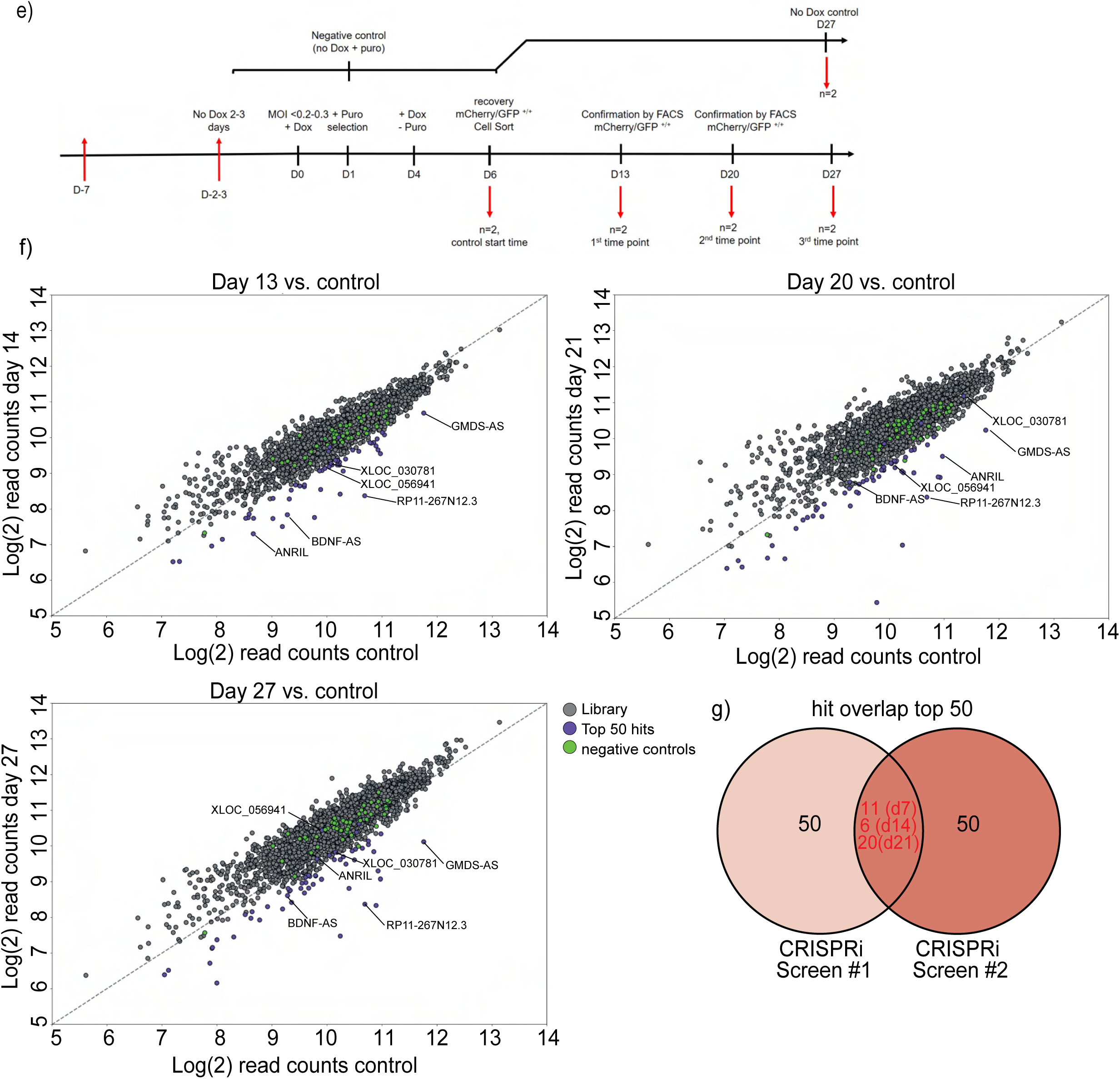
LncRNA CRISPRi screen in melanoma 501mel cells: **a)** Histogram shown raw reads of sgRNA frequency distribution of plasmid library. D8 and D12 refers to selected single cell clone of 501mel-dCas9-KRAB and D7, D14 and D21 for the days of screening time. **b)** Box-plot of log_2_ cpm sgRNA counts during CRISPRi screen. **c)** heatmap showing unsupervised hierarchical clustering of z-score normalized sgRNA expression throughout all samples indicated. **d)** Density dot-plots for lncRNA CRISPRi screen over replicates for correlation (n=2), correlation coefficient is indicated. **e)** time-line of lncRNA CRISPRi screen 2.0. **f)** Dot-plots shown results of CRISPRi lncRNA library screen 22.0 at time points day 7, 14 and 21 post-selection as log2 read counts of control day 0 vs. respective time points. Each dot represents a single sgRNA (?). Selected lncRNA hits from CRISPRi screen as well as lncRNA positive control ANRIL are highlighted in purple as depleted. Negative control sgRNAs are shown in green. **g)** Venn-diagram showing the overlap of lncRNA top 50 hits in both screens at all time points.

**Figure S4:**
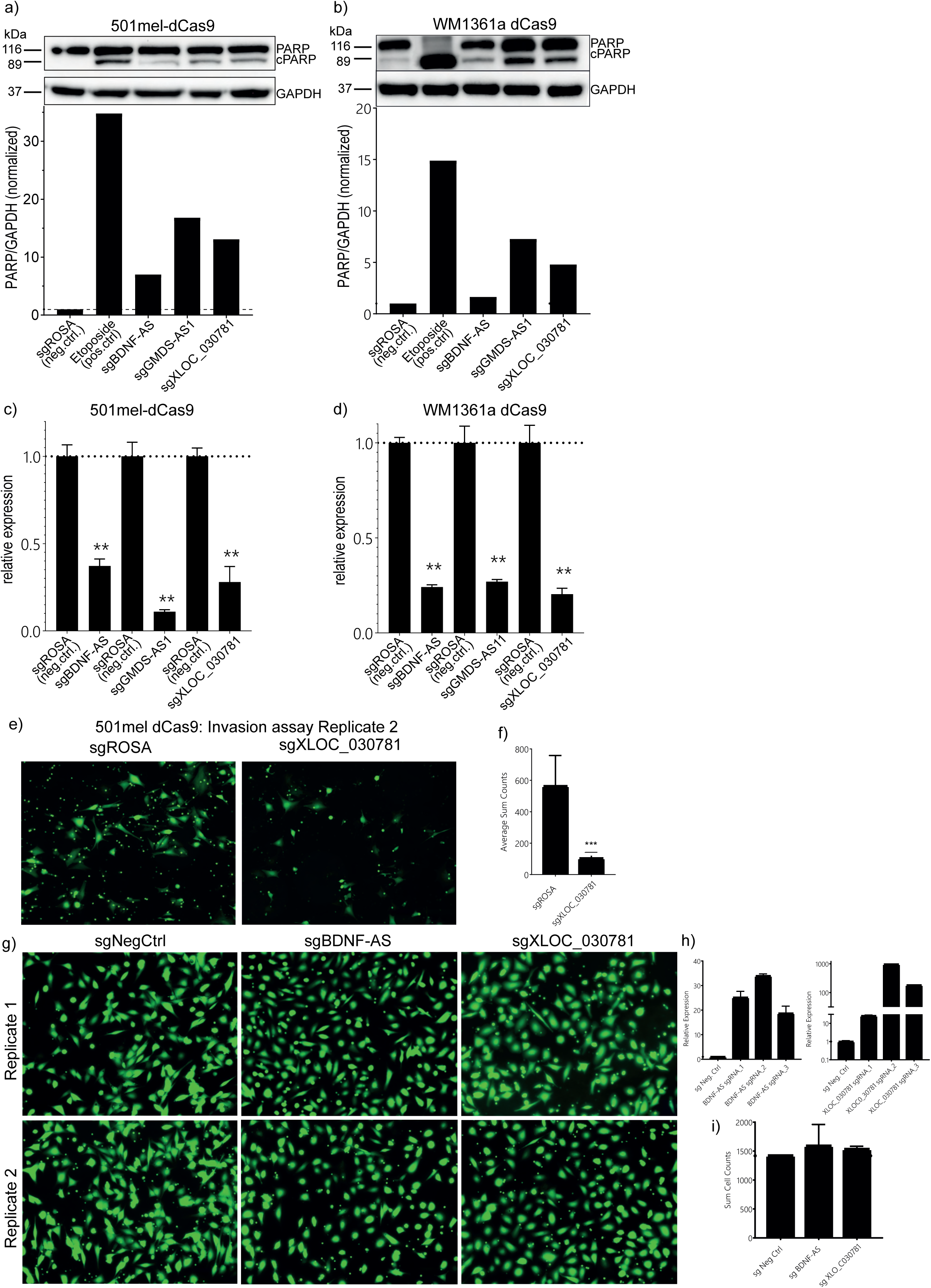
**a)** and **b)** PARP cleavage apoptosis assay replicate 2 by Western-blot and densitometric quantification in 501mel-dCas9-KRAB and WM1361a-dCas9-KRAB (related to Figure 3c). **c)** and **d)** qPCR knockdown confirmation of sgBDNF-AS, GMDS-AS1 and XLOC_030781 in 501mel-dCas9-KRAB and WM1361a-dCas9-KRAB relative to GAPDH and sgROSA control. **e)** Trans-well cell migration assay. Fluorescence photomicrographs show calcein stained migrated cells of sgXLOC_030781 knocked-down 501mel-dCas9-KRAB cells vs. sgROSA negative control and quantification of average sum count cell numbers (replicate 2, related to Figure 3 d) **g-i)** Trans-well cell migration assay. Fluorescence photomicrographs show calcein stained migrated cells of sgXLOC_030781 and sgBDNF-AS 501mel CRISPR-activation cell cells vs. sgNegative control. **h)** confirmation of lncRNA sgXLOC_030781 and sgBDNF-AS overexpression by qPCR and **i)** quantification of average sum count cell numbers.

**Figure S5:**
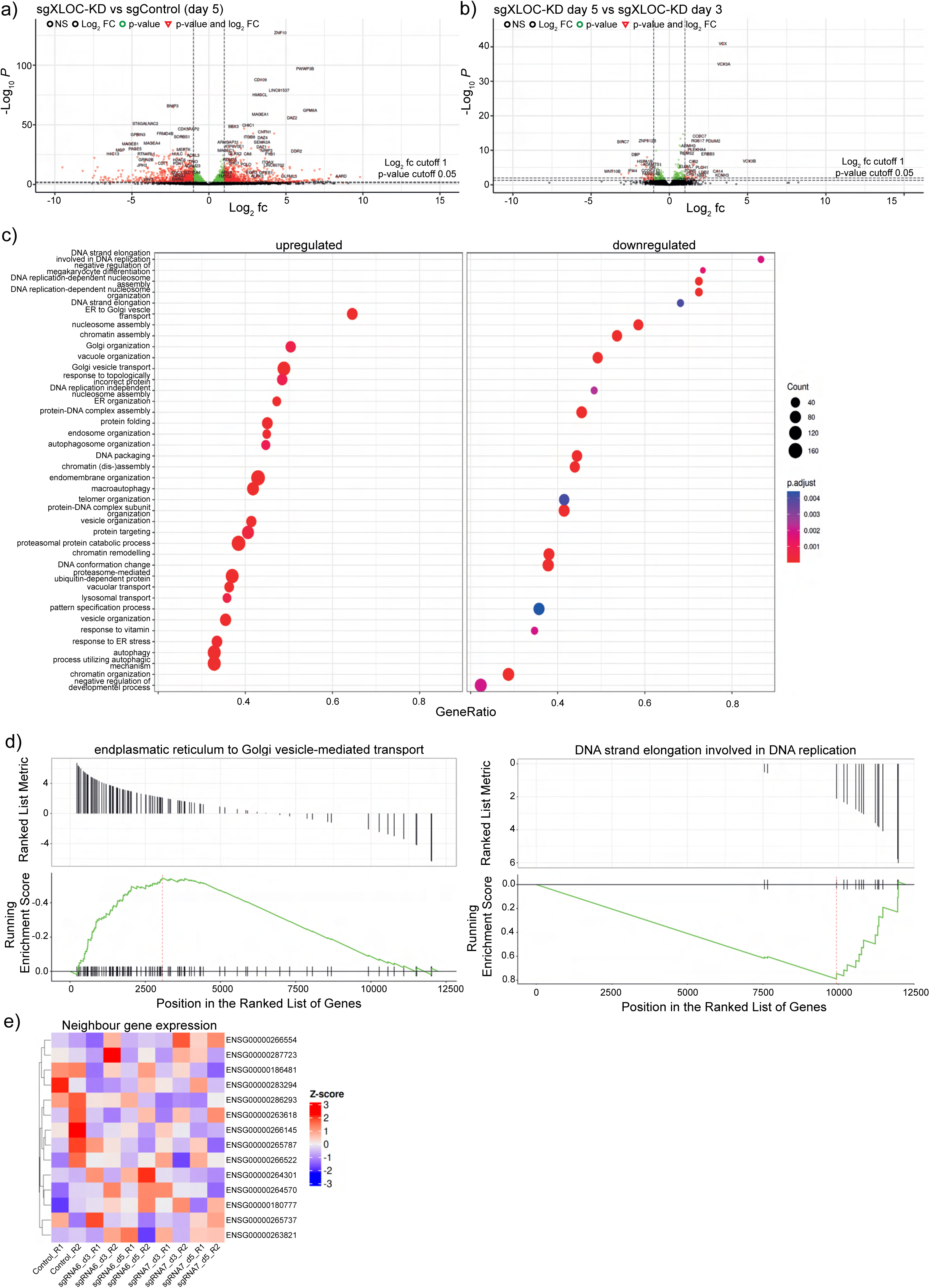
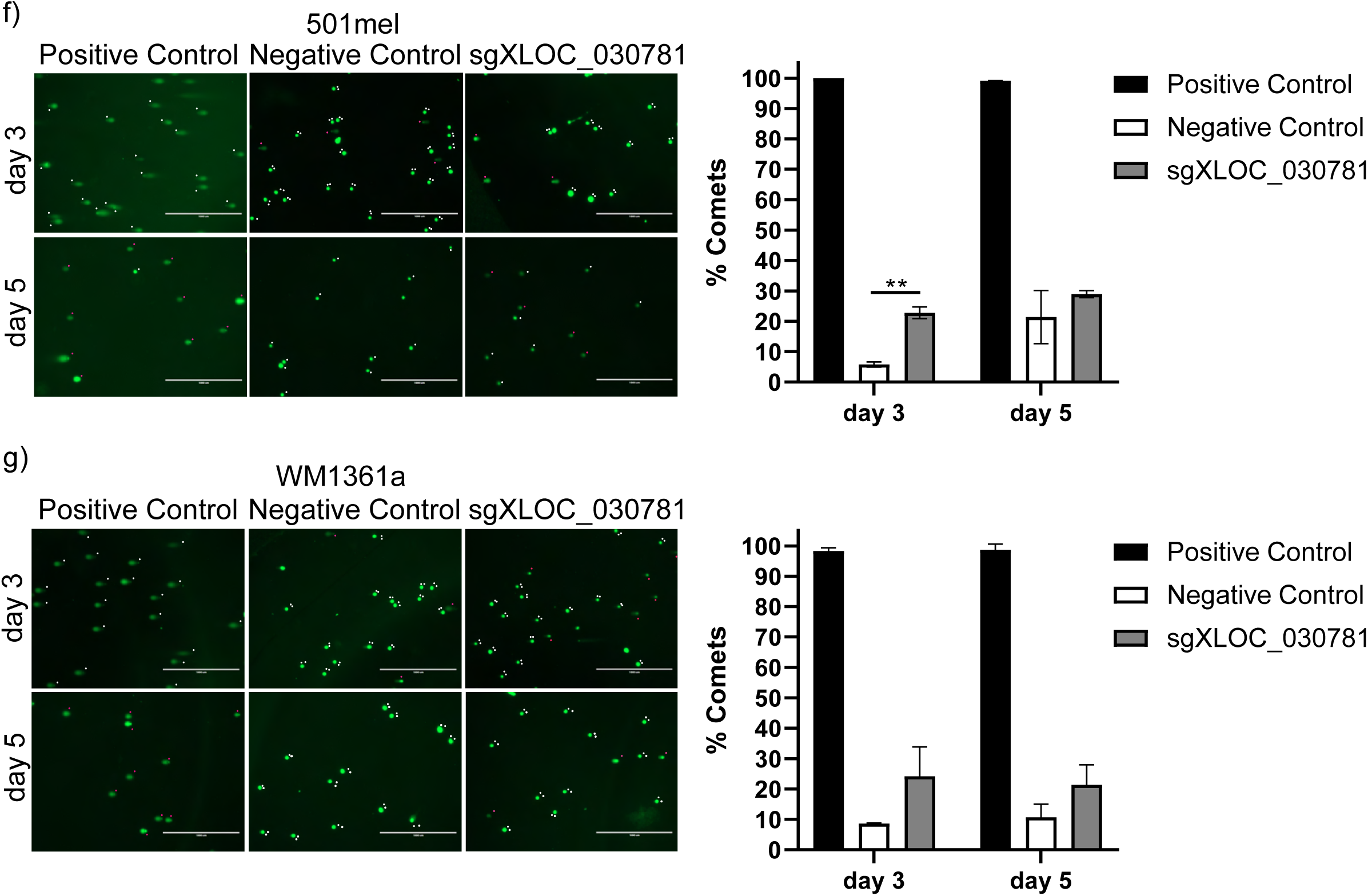
XLOC_030781 CRISPRi knockdown and RNA-sequencing transcriptomic expression analysis in 501mel-dCas9-KRAB at day 5 and additional controls. **a)** Volcano plot of significantly differentially expressed genes (red) upon sgXLOC_030781 knockdown at day 5, p-value cut-off 0.05, log_2_ fc= 1. **b)** Volcano plot expression comparison of sgXLOC_030781 knockdown of day 3 vs. day 5. **c)** Gene ontology enrichment analysis of up- and down regulated gene sets upon sgXLOC_030781 knockdown at day 5. **d)** GSEA enrichment analysis of each top example of up-regulated (ER to Golgi vesicle transport, left) and down-regulated genes (DNA-strand elongation involved in DNA replication, right). **e)** Heatmap shown no significant neighbor gene expression correlation upon sgXLOC_030781 in 501mel-dCas9-KRAB at both time points day 3 and 5. **f)** COMET assay for involvement of XLOC_030781 knockdown in DNA double strand break at day 3 and 5 post-infection in 501mel-dCas9-KRAB and WM1361a-dCas9-KRAB. Right panel shows COMET quantification of sgXLOC_030781 knockdown compared to positive and negative control (see methods for further detail, n=2).

**Figure S6:**
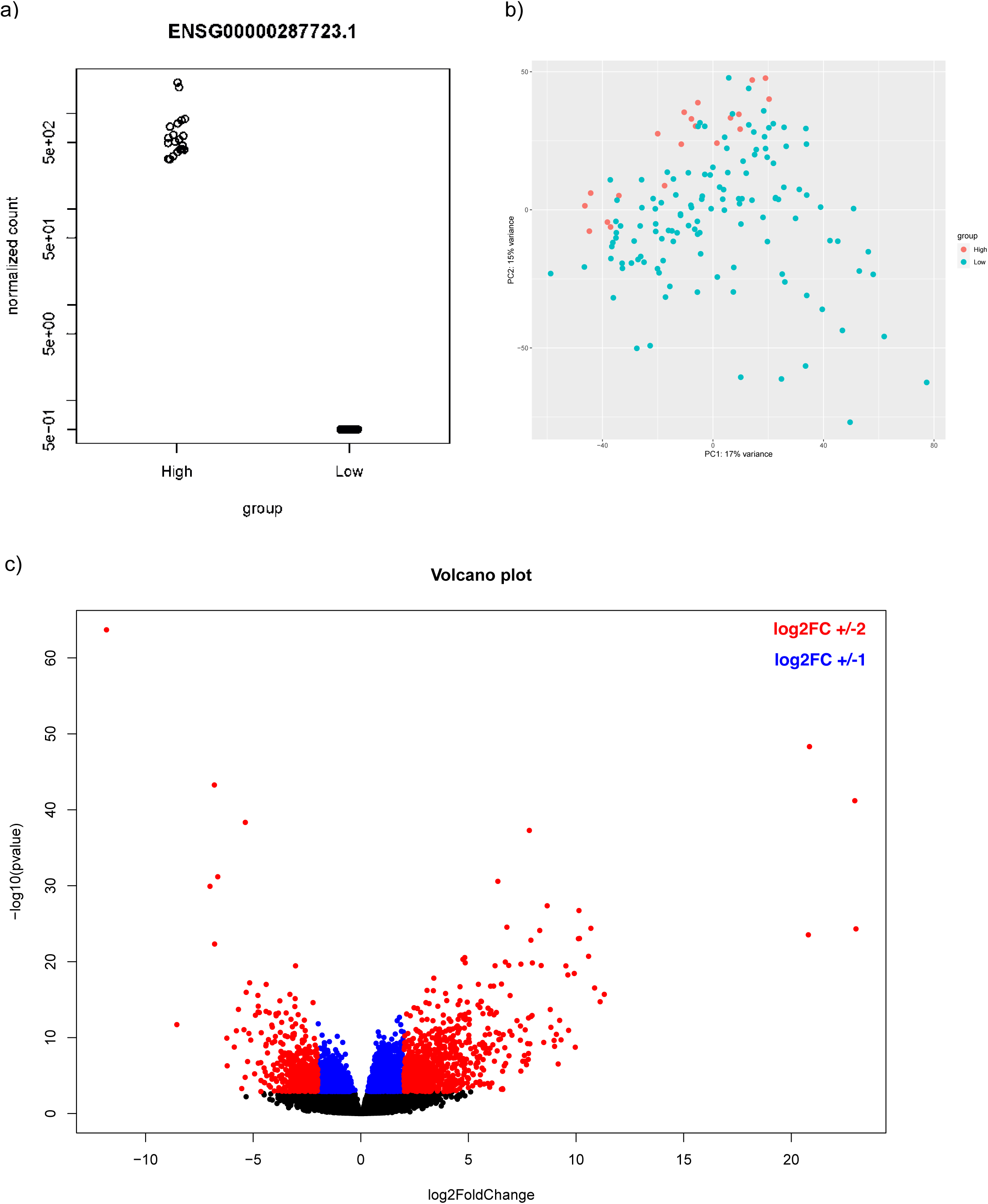
XLOC_030781/ENSG0000287723 lncRNA expression metanalysis in clinical melanoma TCGA samples by DESeq2. **a)** Stratification of 20 melanoma with very high vs 125 patients with absent XLOC_030781/ENSG0000287723 expression. **b)** PCA analysis of high vs. patients with absent XLOC_030781/ENSG0000287723 expression. **c)** Volcano plot of differentially expressed genes in high vs. patients with absent XLOC_030781/ENSG0000287723 expression.

